# Enabling constrained spherical deconvolution and diffusional variance decomposition with tensor-valued diffusion MRI

**DOI:** 10.1101/2021.04.07.438845

**Authors:** Philippe Karan, Alexis Reymbaut, Guillaume Gilbert, Maxime Descoteaux

## Abstract

Diffusion tensor imaging (DTI) is widely used to extract valuable tissue measurements and white matter (WM) fiber orientations, even though its lack of specificity is now well-known, especially for WM fiber crossings. Models such as constrained spherical deconvolution (CSD) take advantage of high angular resolution diffusion imaging (HARDI) data to compute fiber orientation distribution functions (fODF) and tackle the orientational part of the DTI limitations. Furthermore, the recent introduction of tensor-valued diffusion MRI allows for diffusional variance decomposition (DIVIDE), opening the door to the computation of measures more specific to microstructure than DTI measures, such as microscopic fractional anisotropy (μFA). However, tensor-valued diffusion MRI data is not mathematically compatible with latest versions of CSD and the impacts of such atypical data on fODF reconstruction with CSD are yet to be studied. In this work, we lay down the mathematical and computational foundations of a tensor-valued CSD and use simulated data to explore the effects of various combinations of diffusion encodings on the angular resolution of extracted fOFDs. We also compare the combinations with regards to their performance at producing accurate and precise μFA with DIVIDE, and present an optimised protocol for both methods. We show that our proposed protocol enables the reconstruction of both fODFs and μFA on *in vivo* data.

## 1. Introduction

Diffusion MRI (dMRI) [Le Bihan and Breton, 1985] allows for non-invasive probing of the diffusion of water molecules in tissues such as the human brain. In particular, diffusion tensor imaging (DTI) [Basser et al., 1994] models the average voxel content with a diffusion tensor to get access to valuable information about the intra-voxel diffusion profile. Indeed, the diffusion tensor gives insight into the orientation of the white matter (WM) fibers and leads to the calculation of DTI measures, such as the well known mean diffusivity (MD), fractional anisotropy (FA) [Basser and Pierpaoli, 1996], axial diffusivity (AD) and radial diffusivity (RD). While the orientation of the diffusion tensor and the computation of these measures are great tools for the study of WM in the brain and are widely used, important limitations were pointed out [Wheeler-Kingshott and Cercignani, 2009; Jones and Cercignani, 2010]. Furthermore, Tuch et al. [2002] showed that DTI cannot properly model a voxel containing multiple WM fiber orientations. This leads to the FA drop in WM fiber crossings and to a counter-intuitive FA increase in, e.g., Alzheimer lesions [Douaud et al., 2011; Teipel et al., 2014], resulting in a very ambiguous interpretation of this measure in what is estimated to represent 60 to 90% of voxels in a typical whole-brain scan [Descoteaux, 2008; Jeurissen et al., 2013; Volz et al., 2018]. To obtain a better WM fiber orientation profile, Tuch et al. [2002] proposed the high angular resolution diffusion imaging (HARDI) idea, which gave birth to many HARDI-based methods such as constrained spherical deconvolution (CSD) [Tournier et al., 2004, 2007; Descoteaux et al., 2009], used to extract a fiber orientation distribution function (fODF) from HARDI data. These fODFs can then serve as guides for tractography algorithms [Mori et al., 1999; Poupon et al., 2000; Catani et al., 2002], allowing structural connectivity human brain mapping studies and applications.

An extension of DTI, diffusion kurtosis imaging (DKI) [Jensen et al., 2005], enables to estimate a first measure of tissue heterogeneity: the mean kurtosis. However, this mean kurtosis is not specific, as it originates from both microscopic anisotropy (probing pure cell elongation) and isotropic heterogeneity (variance of isotropic diffusivities or variance of cell densities). A new dMRI technique introduced in the mid-2010s, called tensor-valued dMRI [Eriksson et al., 2013, 2015; Westin et al., 2014, 2016] or b-tensor encoding, shows great promise in alleviating the lack of specificity of conventional dMRI techniques such as DTI or DKI. In particular, Lasič et al. [2014] proposed a way to disentangle the isotropic and anisotropic components of the diffusional variance, leading to new measures of microscopic anisotropy and isotropic heterogeneity. Recent papers [Nilsson et al., 2020; Naranjo et al., 2021] have even established clinically feasible tensor-valued dMRI scans providing sufficient data to compute these new measures with similar methods. Moreover, many studies have investigated the potential of tensor-valued diffusion encoding for microstructural characterizations of brain tumors and neurodegenerative diseases [Szczepankiewicz et al., 2015, 2016; Andersen et al., 2020; Kamiya et al., 2020; Nilsson et al., 2020].

Several studies have accounted for sub-voxel WM fascicle orientations while employing tensor-valued diffusion encoding to capture the aforementioned diffusion measures [Cottaar et al., 2020; Reymbaut et al., 2020a,b, 2021; de Almeida Martins et al., 2021]. However, much remains to be done in evaluating the effects of various diffusion encodings on fODF reconstruction with CSD. Indeed, the standard single-shell single-tissue CSD (ssst-CSD) [Tournier et al., 2007; Descoteaux et al., 2009] does not allow for the use of tensor-valued dMRI data and the current state of the literature does not provide any CSD model that can use such data as input, except for one conference abstract [Jeurissen and Szczepankiewicz, June 2018]. Nevertheless, the work of Jeurissen et al. [2014] on extending the ssst-CSD model to a multi-shell multi-tissue CSD (msmt-CSD) model is a great example of the flexibility of CSD, allowing it to take multi-shell dMRI data as input and accurately differentiate brain tissues.

This paper establishes the foundations of a tensor-valued diffusion encoding CSD model, as a mathematical extension of msmt-CSD, enabling the reconstruction of fODFs using tensor-valued dMRI data. The impacts of different combinations of b-tensor shapes, namely linear, planar and spherical tensors, on the reconstruction of fODFs are explored using the adapted CSD model on simulated data. In parallel, these combinations are also challenged with a method for disentangling the diffusional variance, and performances on both methods are compared to lead to the overall best combination. It is important to propose a technique that can extract both accurate crossing fibers as well as advanced microstructural maps from tensor-valued dMRI. Thus, we propose a 10 minutes long tensor-valued dMRI protocol enabling an accurate reconstruction of the fODFs while also allowing the computation of the b-tensor encoding microstructure measures. The performances of this final protocol are shown on *in vivo* data as a demonstration of the new tensor-valued diffusion encoding CSD model’s potential.

## 2. Theory

Assuming a non diffusion-weighted (DW) signal *S*_0_, the DW signal *S_i_* for an acquisition direction *i* is defined as

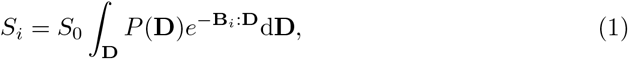

where *P*(**D**) is the intra-voxel diffusion tensor distribution (DTD) and **B**_*i*_ : **D** denotes the Frobenius inner product between the so-called b-tensor **B**_*i*_ and a diffusion tensor **D**:

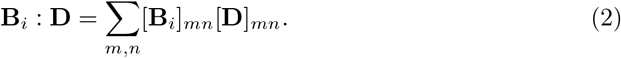

Note that this definition of the signal assumes a heterogeneous intra-voxel content, described by the DTD [Basser and Pajevic, 2003; Jian et al., 2007; Reymbaut, 2020].

### 2.1. Diffusional variance from the DTD

Following the formalism of Szczepankiewicz [2016], different measures can be calculated from the DTD *P*(**D**). For instance, the average (denoted by ⟨·⟩) of the diffusion tensors across the DTD in a voxel gives a voxel-scale diffusion tensor ⟨**D**⟩, the same one that is at the center of the DTI model. From this, it is well known that the mean diffusivity (MD) is obtained according to MD= E_λ_[⟨**D**⟩], where E_λ_[·] is the average over tensor eigenvalues λ. However, the MD can also be calculated as the average of all isotropic diffusivities E_λ_[**D**] in the DTD:

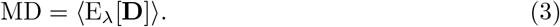

The DTD also contains information about the isotropic variance *V*_I_, reflecting the isotropic heterogeneity of the voxel, and the anisotropic variance *V*_A_, describing the microscopic anisotropy of the voxel. These diffusional variances are calculated as follow, with V_λ_[·] being the population variance of tensor eigenvalues λ:

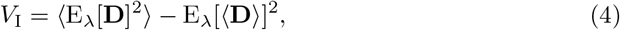

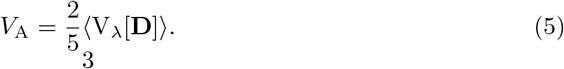

Diffusional variance decomposition (DIVIDE) can be used to extract MD, *V*_I_ and *V*_A_ from tensor-valued dMRI data with at least two different b-tensor encodings Lasič et al. [2014]; Szczepankiewicz et al. [2015, 2016]. This method estimates the isotropic and anisotropic variances of the DTD by fitting the following inverse Laplace transform of the gamma distribution function [Röding et al., 2012] to powder-averaged tensor-valued dMRI data 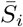:

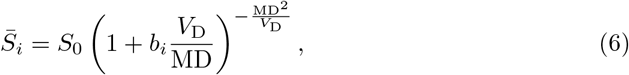

where 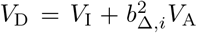 is the diffusional variance and *b*_Δ*,i*_ is a factor describing the encoding shape (*b*_Δ*,i*_ = 1 for linear, *b*_Δ*,i*_ = − 0.5 for planar and *b*_Δ*,i*_ = 0 for spherical). This disentanglement of the diffusional variances enables the definition of the microscopic fractional anisotropy μFA [Lasič et al., 2014; Reymbaut, 2020] as:

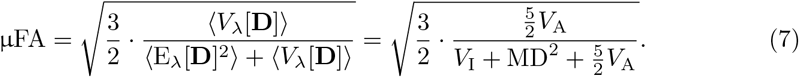

This new measure is thus representative of the average anisotropy computed across all microscopic environments in a voxel, with values going from 0 (purely isotropic cellular structures) to 1 (purely stick-like cellular structures). In comparison, the typical FA computed from DTI gives information about the anisotropy observed at the voxel scale, consequently depending on the orientation coherence of the underlying cellular structures, which is quantified by the order parameter (OP) [Lasič et al., 2014]. The μFA becomes equal to the FA in voxels where all microscopic environments are perfectly identical and ordered.

### 2.2. Tensor-valued constrained spherical deconvolution

In the case of a single homogeneous axisymmetric fiber with orientation **u**_*k*_ ≡ (*θ_k_, ϕ_k_*), the DTD from equation 1 becomes a Dirac distribution peaked at a single diffusion tensor **D**_*k*_ of main eigenvector **u**_*k*_, axial and radial diffusivities *D*_||_ and *D*_⊥_. The Frobenius inner product from equation 2 then becomes

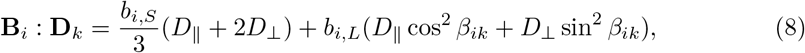

where the angle *β_ik_* separating the orientation of the b-tensor **e**_*i*_ ≡ (Θ_*i*_, Φ_*i*_) and the fiber orientation **u**_*k*_ is given by the spherical law of cosines

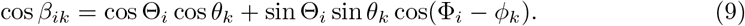

The couple (*b_i,S_, b_i,L_*) determines the b-tensor’s encoding type such as

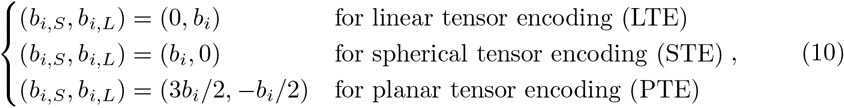

where *b_i_* is the typical b-value, given by *b_i_* = Tr(**B**_*i*_), the trace of the b-tensor.

With the DTD being a Dirac distribution peaked at a single diffusion tensor and substituting the Frobenius inner product from equation 8, equation 1 simplifies to

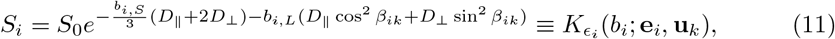

where we introduce the single fiber response function (FRF) or convolution kernel 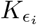, which changes according to the diffusion encoding of acquisition *i*, such that *ϵ_i_* ∈ {linear, planar, spherical}. If multiple identical homogeneous axisymmetric fibers are present, the equation becomes an integral over the unit sphere 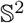

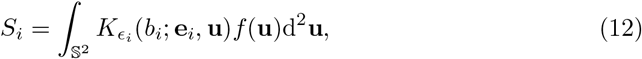

where the diffusion signal is modeled as the convolution of an fODF *f* (**u**) with a fiber response function 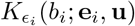. The fODF weighs different unit orientations of fiber in the signal, while the response function corresponds to the DW signal of a single fiber with orientation **u**.

Using equation 10, the fiber response functions for linear, planar and spherical tensor encoding are

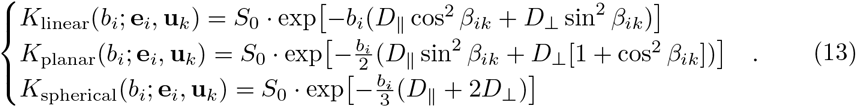

Figure 1 shows these encoding dependent theoretical FRFs for multiple b-values and tissues, namely white matter, grey matter (GM) and the cerebrospinal fluid (CSF), from diffusivity values taken from the literature and real data examples (see section 3.4).

**Figure 1:**
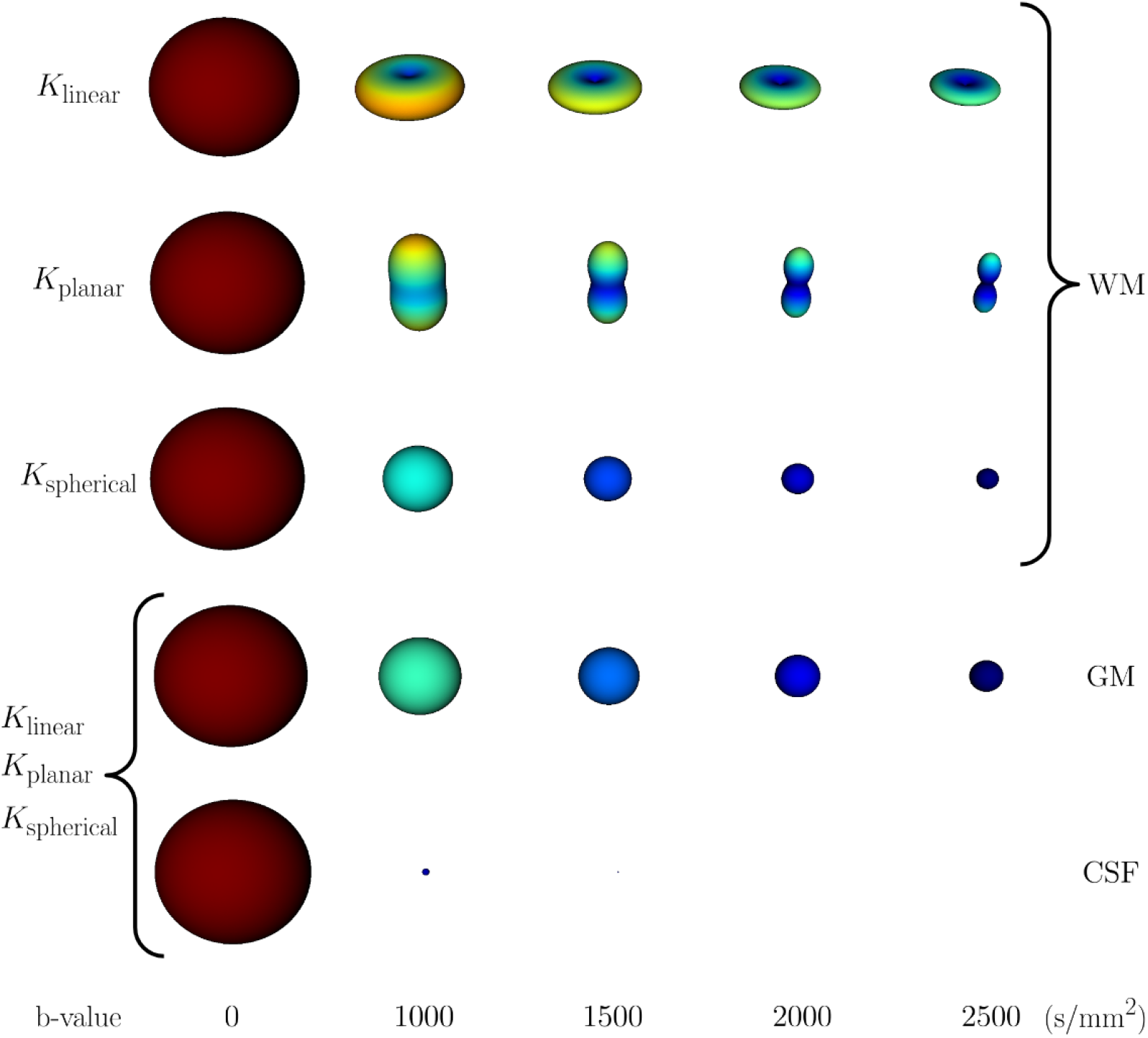
Visualisation of fiber response functions (FRF) 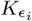 computed from equation 13 for different b-values and tissues. The WM tissue, defined by *D*_||_ = 1.7 *×* 10^−3^mm^2^*/*s and *D*_⊥_ = 0.3 *×* 10^−3^mm^2^*/*s, shows varying FRFs for LTE, PTE and STE. For GM (*D*_||_ = *D*_⊥_ = 0.6 *×* 10^−3^mm^2^*/*s) and CSF (*D*_||_ = *D*_⊥_ = 3.0 *×* 10^−3^mm^2^*/*s), all tensor encodings produce the same FRF. Every fiber response function is calculated using *S*_0_ = 1 and a diffusion tensor pointing in the *z*-axis. The FRFs for CSF are too small to be visible at *b* ≥ 2000 s/mm^2^, because the diffusivities are high and lead to the amplitude of each *b* > 1000 s/mm^2^ losing a factor of approximately 4.5 from the amplitude of the previous shell.

The fODF is usually expressed as a linear combination of *N*_SH_ basis functions *Y_j_* (**u**), such as spherical harmonics (SH), leading to

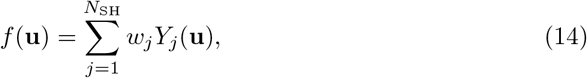

where *w_j_* are the SH coefficients in the case of an SH basis. Applying this to equation 12 then gives

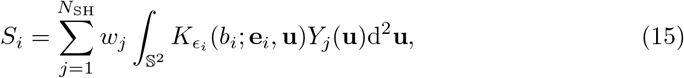

which can be written simply as a linear problem:

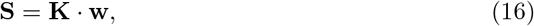

where the following matrix **K** and column vectors **S** and **w** read:

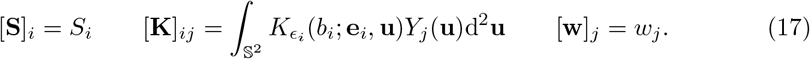

Note that in the case of a linear encoding only acquisition, equation 15 falls back to the classic single-shell or multi-shell formulation, with a single linear fiber response function per b-value. The linearized problem of equation 16 can then be solved using the msmt-CSD model developed by Jeurissen et al. [2014]. In this paper, the authors expand the ssst-CSD method from Tournier et al. [2007]; Descoteaux et al. [2009] to allow *m* shells and *n* tissues, resulting in the following constrained linear least squares problem:

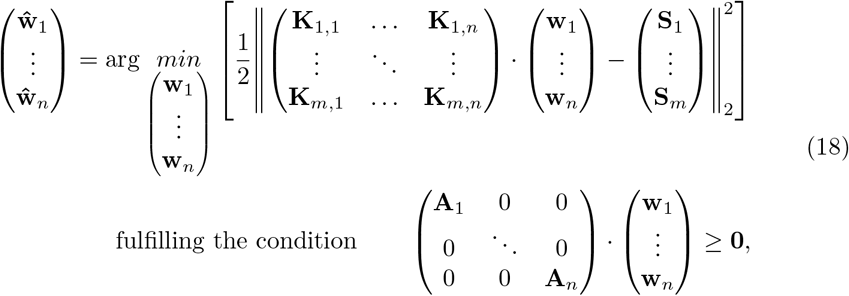

where **A**_*j*_ is the matrix relating the coefficients **w**_*j*_ to the fODF amplitudes. Solving this equation leads to the unknown vectors of coefficients of the fODF, with one vector for each tissue. In the case of *m* = *n* = 1, this equation goes back to the ssst-CSD formalism.

## 3. Methods

### 3.1. Implementation of tensor-valued constrained spherical deconvolution

In the case of tensor-valued dMRI data, the linearized problem of equation 16 can be solved using an extension of equation 18 that enables CSD with multiple tensor-valued encodings, a method we call multi-encoding msmt-CSD (memsmt-CSD). This allows for *m_E_* shells per encoding *ϵ*, and *n* tissues. The convolution kernels then become a concatenation of the different encoding kernels, such as the linear (**K**^L^ with *m*_L_ shells), planar (**K**^P^ with *m*_P_ shells) and spherical (**K**^S^ with *m*_S_ shells) ones. The signals vector becomes a concatenation of the signal vectors from different encodings, such as the LTE (**S**^L^), the PTE (**S**^P^) and the STE (**S**^S^) ones, leading to the following constrained linear least squares problem:

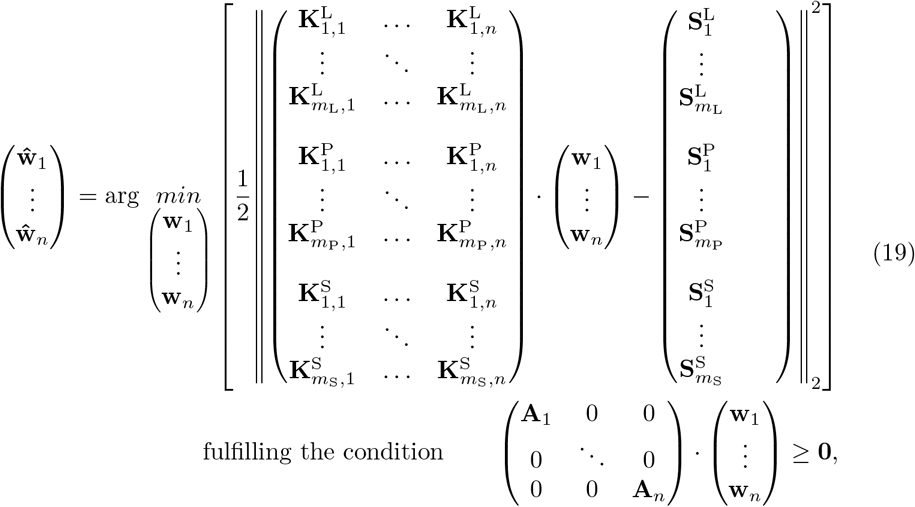

where **A**_*j*_ is the matrix relating the coefficients **w**_*j*_ to the fODF amplitudes. Again, solving this equation leads to the unknown vector of fODFs coefficients for each tissue. From these, the volume fraction (VF) of each tissue can be calculated as the amplitude of their first coefficient, leading to a sort of tissue classification.

Both constrained linear least squares problems from equations 18 and 19 can be rewritten as a strictly convex quadratic programming (QP) problem, using the compact nomenclature of equation 16 such that:

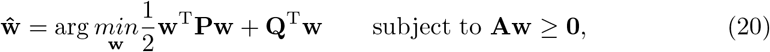

where **P** = **K**^T^**K** and **Q** = −**K**^T^**S**. This QP problem is solved with DIPY [Garyfallidis et al., 2014], which uses a QP solver implemented in CVXPY [Diamond and Boyd, 2016; Agrawal et al., 2018], based on the OSQP solver [Stellato et al., 2020]. The fiber response functions 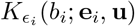 for each encoding and b-value pair are calculated from equation 13 using diffusivities extracted from a DTI fit for each tissue. Figure 2 summarizes the computational steps of tensor-valued constrained spherical deconvolution.

**Figure 2:**
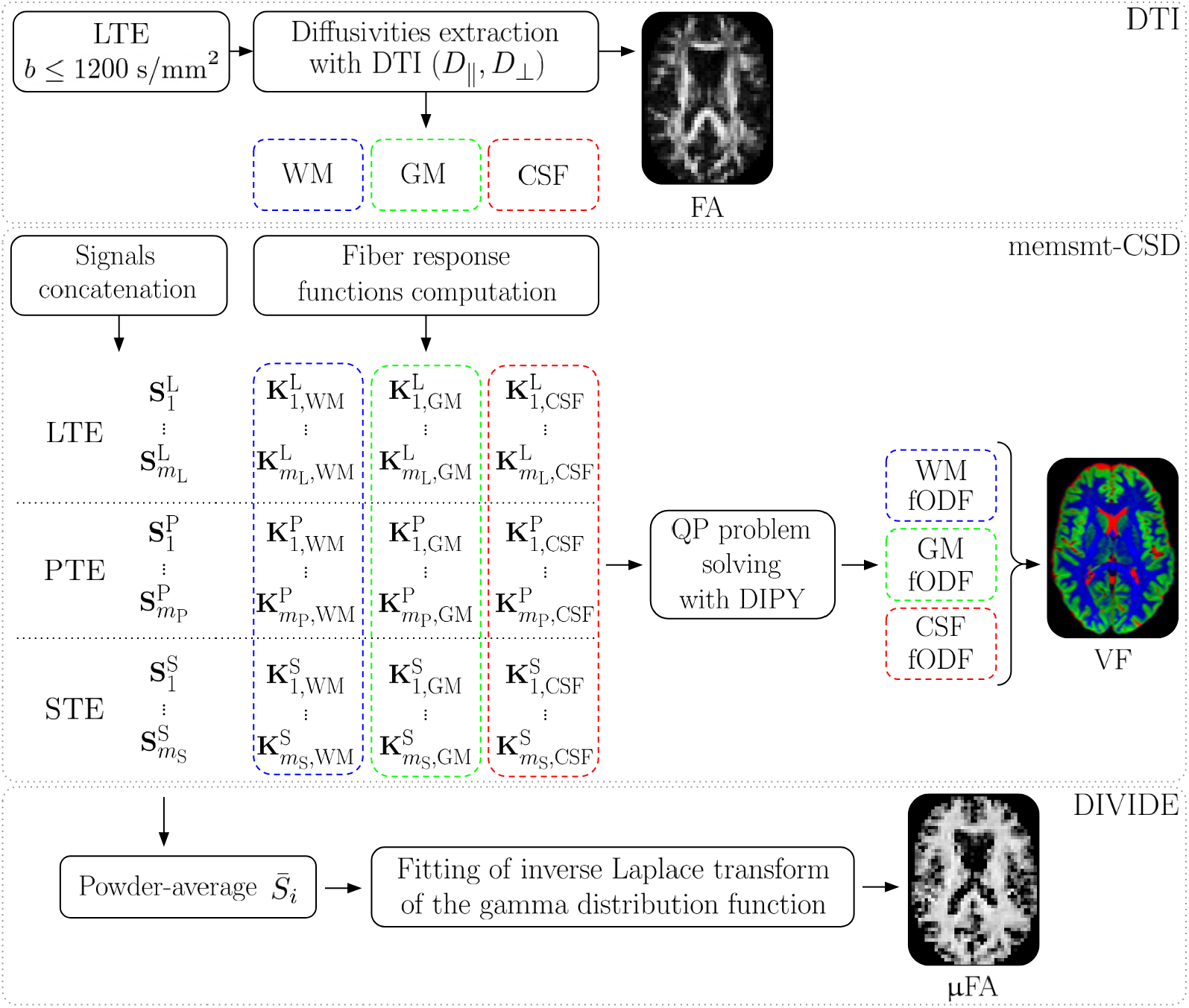
Summary of the computational steps used in this study, comprising DTI, memsmt-CSD and DIVIDE, for linear, planar and spherical tensor encodings, and three tissues (WM, GM and CSF). FA and μFA images are adapted from Szczepankiewicz et al. [2015].

### 3.2. Measures computation

The DTI measures such as FA and diffusivities (AD, MD, RD) are calculated using the eigenvalues of the diffusion tensor obtained from a DTI fit, as shown by Basser and Pierpaoli [1996]. The fit is performed only on the linear encoding data and with shells below *b* = 1200 s/mm^2^.

The measures enabled by the combination of at least two different b-tensor encodings are obtained using the DIVIDE method. The implementation of this method is strongly inspired by Nilsson et al. [2018]. Note that although DIVIDE relies on powder-averaged signals, it has suggested to be, to some extent, robust to not perfectly rotation invariant signals [Reymbaut et al., 2020c]. The present study focuses on the microscopic fractional anisotropy (μFA) introduced by Lasič et al. [2014] and calculated using equation 7, in comparison to the FA. Figure 2 summarizes the processing steps leading to the FA and μFA.

### 3.3. Simulating tensor-valued diffusion data

To study the impact of different combination of diffusion encoding shapes and number of directions per shells on the fODFs and the computation of the μFA, tensor-valued diffusion data is simulated using a discrete version of equation 1. The first step is to choose a set of *N_k_* diffusion tensors **D**_*k*_ that will describe the composition of the voxel, each of them being associated with a tissue compartment. These diffusion tensors are defined by their eigenvalues in their principal axis system (PAS), such that

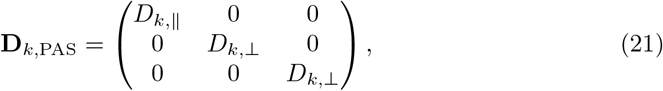

where *D_k,_*_||_ and *D_k,_*_⊥_ are the axial and radial diffusivities, respectively. For each of these diffusion tensors, *D_k,_*_||_ and *D_k,_*_⊥_ are converted into the isotropic diffusivity *D_k,_*_iso_ and the normalized anisotropy *D_k,_*_Δ_ [Conturo et al., 1996], describing the anisotropy of the diffusion tensor, following the relations *D_k,_*_iso_ = (*D_k,_*_||_ + 2*D_k,_*_⊥_)*/*3 and *D_k,_*_Δ_ = (*D_k,_*_||_ − *D_k,_*_⊥_)*/*(3*D_k,_*_iso_). Then, both *D_k,_*_iso_ and *D_k,_*_Δ_ become the mean of discrete Gaussian distributions with a given relative standard deviation (STD) *σ_k_* (relative to its mean) and *N* discrete elements. These two Gaussian distributions are weighted by the volume fraction and the non-DW signal 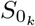 of the tissue compartment they represent. The distribution centered at *D_k,_*_iso_ is also flipped to ensure a negative covariance between the two distributions. Then, the distributions of *D_k,_*_iso_ and *D_k,_*_Δ_ are converted back to *D_k,_*_||_ and *D_k,_*_⊥_ distributions, creating a distribution of diffusion tensors 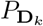 described by the distributions of axial and radial diffusivities. Every diffusion tensor in the DTD is rotated by given angles *θ_k_* and *ϕ_k_*, corresponding to the orientation of their initial mean diffusion tensor **D**_*k*_. The angles *θ_k_* and *ϕ_k_* follow the physics convention for spherical coordinates, meaning that *θ_k_* gives the angle with respect to the *z* axis, while *ϕ_k_* gives the angle with respect to the *x* axis. Finally, the *N_k_* weighted DTD are concatenated into a single DTD 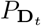 representing the whole voxel’s content with *N_t_* = *N_k_* · *N* diffusion tensors **D**_*t*_. This allows the computation, for any chosen b-tensor **B**_*i*_, of the diffusion signal in the voxel:

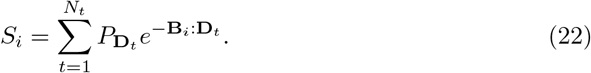

The signal obtained from this process can also be subject to added noise, which is calculated with respect to a given *S*_0_ and signal to noise ratio (SNR), relative the chosen *S*_0_:

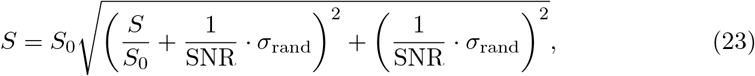

where *σ*_rand_ is a random number going from 0 to 1.

From the DTD, the ground truth μFA is calculated using equations from section 2.1.

### 3.4. Simulated anatomy

The simulation method previously described is used to generate the DW signals of five typical voxels, representing the fictional anatomy of this study. These voxels are composed of various volume fractions of four different tissue compartments (*N_k_* = 4), each compartment *k* being defined by the parameters *D_k,_*_||_, *D_k,_*_⊥_, *θ_k_*, *ϕ_k_*, *σ_k_* and *S*_0*,k*_, with *N* = 100 for each of them. Table 1 shows the parameters configuration for each tissue compartment, comprising two identical WM compartments separated by a certain angle *α*, one grey matter (GM) compartment and one cerebrospinal fluid (CSF) compartment. Table 2 presents the composition of the five simulated voxels. The first voxel is a WM fiber crossing of equal proportions, described by the separation angle *α*. The second voxel is a single WM fiber, while the fourth and fifth voxels are 100% GM and CSF, respectively. The third voxel represents a partial volume between voxel 2 and voxel 4, composed of 50% WM fiber and 50% GM.

**Table 1:**
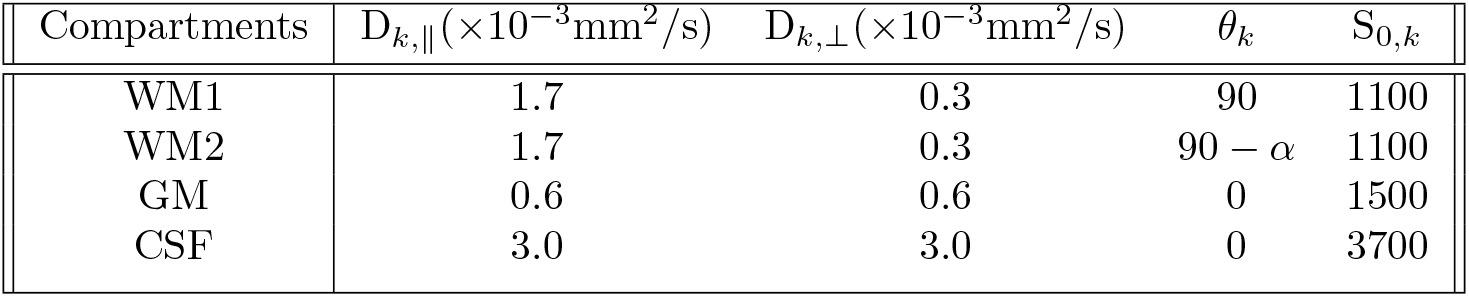
Parameters configuration of the four tissue compartments used to generate simulated data. The two WM compartments are built from the same diffusivities and only differ by the orientation of their diffusion tensor **D**_*k*_, separated by *α* degrees in the *y*-*z* plane. The angle *θ_k_* for GM and CSF are both equal to 0 since their diffusion tensor is isotropic, thus not depending on orientation. For all compartments, *ϕ_k_* = 0 and *σ_k_* = 0.15. A different non-DW signal *S*_0*,k*_ is set for each tissue type.

**Table 2:** Voxels composition, described by the volume fractions, in percentage, of each tissue compartment.

| Compartments | Voxel 1 | Voxel 2 | Voxel 3 | Voxel 4 | Voxel 5 |
| --- | --- | --- | --- | --- | --- |
| WM1 | 50 | 100 | 50 | 0 | 0 |
| WM2 | 50 | 0 | 0 | 0 | 0 |
| GM | 0 | 0 | 50 | 100 | 0 |
| CSF | 0 | 0 | 0 | 0 | 100 |

The choice of the axial and radial diffusivities is motivated by typical *in vivo* diffusivity values. WM and CSF diffusivities are inspired by Pierpaoli and Basser [1996]; Alexander [2008]; Alexander et al. [2010]; Zhang et al. [2012], while GM diffusivity value comes from Pierpaoli and Basser [1996]; Liu et al. [2006]. Moreover, the non-DW signal *S*_0*,k*_ set for WM, GM and CSF are approximated from data obtained from the MGH-USC Human Connectome Project database [Glasser et al., 2013; Sotiropoulos et al., 2013].

Throughout the simulation experiments, only the angle *α* and the SNR vary, as does the set of b-tensors **B**_*i*_. The relative STD *σ_k_* stays constant at 15%, providing some variance to the distribution of diffusion tensors composing the voxels. To test the limits of memsmt-CSD, the WM fiber crossing of voxel 1 is simulated many times using different values of *α*, from 90 to 50 degrees. The data is simulated without noise (SNR= ∞) and with SNR=30, SNR=20 and SNR=15, inspired from *in vivo* values of data discussed in later sections. These allow to study the effects of noise on the memsmt-CSD and the DIVIDE processes, as well as the effects of spatial resolution and echo time (TE). Indeed, higher spatial resolution or shorter TE can be mimicked by a higher SNR, for example by comparing SNR=20 to SNR=15.

### 3.5. Simulated datasets

The fictional anatomy set up is used to explore and test different acquisition protocols, each represented by a set of b-tensors **B**_*i*_. These schemes can also be described by a set of b-values, a number of diffusion encoding gradient directions and different choices of encoding shapes. This representation is favoured over the b-tensor itself, as it is easier to grasp and compare to conventional acquisition schemes. The chosen b-values and number of gradient directions per shell, adapted from Nilsson et al. [2020] to allow memsmt-CSD, are presented in table 3 as L, P_1_, P_2_, S_1_ and S_2_. A typical multi-shell multi-tissue acquisition [Theaud et al., 2020] is also added as L_msmt_ for comparison purposes, as it contains the same total amount of directions as L and S_2_ combined.

**Table 3:** Six gradient tables chosen for this study. Each column shows the number of encoding directions in the corresponding shell. The last row shows the total number of directions, including *b* = 0 s/mm^2^.

| b-value<br>(s/mm<sup>2</sup>) | Linear<sub>1</sub><br>(L) | Linear<sub>2</sub><br>(L<sub>msmt</sub>) | Planar<sub>1</sub><br>(P<sub>1</sub>) | Planar<sub>2</sub><br>(P<sub>2</sub>) | Spherical<sub>1</sub><br>(S<sub>1</sub>) | Spherical<sub>2</sub><br>(S<sub>2</sub>) |
| --- | --- | --- | --- | --- | --- | --- |
| 0 | 3 | 5 | 1 | 2 | 1 | 2 |
| 100 | 3 | 0 | 3 | 6 | 3 | 6 |
| 700 | 3 | 8 | 3 | 6 | 3 | 6 |
| 1200 | 12 | 30 | 6 | 10 | 6 | 10 |
| 1800 | 18 | 0 | 6 | 16 | 6 | 16 |
| 2400 | 24 | 60 | 0 | 0 | 0 | 0 |
| Total | 63 | 103 | 19 | 40 | 19 | 40 |

The encoding directions are distributed between shells and encodings using the electrostatic repulsion implementation of Caruyer et al. [2013] (with default inter-shell coupling weights). More precisely, the L and L_msmt_ directions are first both calculated separately from the other encodings. Then, the directions are recalculated when L is used in a protocol with planar encoding, to get the best directional coverage over the shells and the encodings. Directional considerations are different for spherical encoding, as the framework laid out in Section 2 presents spherical diffusion encoding as a perfectly isotropic measurement. In practice, anisotropy in the frequency content of spherically encoded gradient waveforms [Lundell and Lasič, 2020; Szczepankiewicz et al., 2020] and eddy currents [Szczepankiewicz et al., 2020] create situations wherein spherical b-tensors should preferably be rotated. This means that there is a set of directions for L when it is alone or with spherical encoding only, and different sets of directions when it is combined with P_1_ or P_2_, which we call L* and L**, respectively. The spherical encoding directions from S_2_ are also separately calculated using the same method, and the directions of S_1_ are subsampled from it. The subsampling of the encoding directions is done using a method that chooses the directions such as to minimize the electrostatic repulsion energy, based on the implementation from Caruyer et al. [2013]. All of these gradient directions distributions are shown and available at figure A.14.

The gradient tables described in table 3 are combined in different ways to test the impact of diffusion encodings and number of encoding directions on both memsmt-CSD and DIVIDE, while varying the *α* angle and the SNR. The tested protocols are presented in table 4. Note that we add a separate protocol, L_msmt_, corresponding to the gradient table L_msmt_ and only tested with memsmt-CSD.

**Table 4:** Eight protocols tested in the study (first row). These are combinations of gradient tables from table 3 (second row). Note that the set of directions for L changes depending on whether or not it is combined with either of the planar encoding gradient tables. These linear encoding directions sets are defined as L (without planar), L* (with P_1_) and L** (with P_2_).

|  |  |  |  |  |  |  |  |
| --- | --- | --- | --- | --- | --- | --- | --- |
| $L$ | $LP_1$ | $LP_2$ | $LS_1$ | $LS_2$ | $LP_1S_1$ | $LP_2S_1$ | $LP_2S_2$ |
| $L$ | $L^*, P_1$ | $L^{**}, P_2$ | $L, S_1$ | $L, S_2$ | $L^*, P_1, S_1$ | $L^{**}, P_2, S_1$ | $L^{**}, P_2, S_2$ |

Each of these protocols are simulated *N*_rep_ = 1000 times, to avoid outliers due to noise. All the outputs of memsmt-CSD, the DTI fit and DIVIDE are averaged over the *N*_rep_ repetitions. Their standard deviation allows to keep track of the variability of the process and compute the precision of it.

### 3.6. In vivo acquisitions

To confirm the conclusions drawn from the simulated data, protocol LS_2_ was acquired *in vivo* on two healthy male volunteers. These *in vivo* data acquisitions were taken on a Philips Ingenia 3T system with a 32-channel head coil (Philips Healthcare, Best, The Netherlands). Tensor-valued dMRI was made possible by a prototype spin-echo single-shot EPI sequence that enables the use of arbitrary b-tensor shapes for diffusion encoding. Planar and spherical encoding were achieved using asymmetric gradient waveforms, optimized to minimize TE using a constrained optimization method described in Sjölund et al. [2015], with the following settings: Euclidian norm, heat dissipation factor 0.7, amplitude limit of 45 mT/m and a slew rate limit of 90 T/m/s. Imaging was performed on the two volunteers at different spatial resolutions, with the following set of constant parameters: SENSE = 2, Multiband-SENSE = 2, and partial-Fourier = 0.65, and two sets of parameters depending on the resolution. The set at 2.5 mm isotropic (first volunteer) was: resolution = 2.5 × 2.5 × 2.5 mm^3^, TE = 117 ms, TR = 5.6 s, FOV = 240 × 240 mm^2^, slices = 48, while the set at 2.0 mm isotropic (second volunteer) was: resolution = 2 × 2 × 2 mm^3^, TE = 119.5 ms, TR = 7.2 s, FOV = 224 × 224 mm^2^, slices = 60. Given these sets of parameters, the acquisition times that would have each protocol from table 4 are presented in table 5. Note that the significant increase of time compared to the 3 minutes protocol of Nilsson et al. [2020] is due to the added number of directions for LTE, the different MRI scanner and the increased resolution for the set at 2 mm isotropic.

**Table 5:**
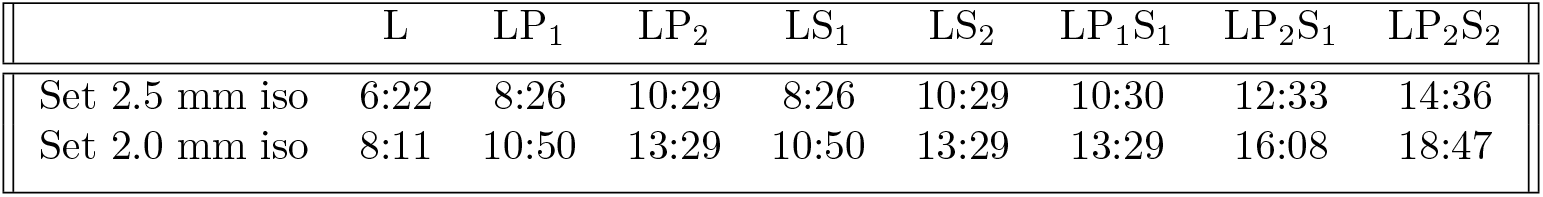
Real acquisition times of the protocols presented in table 4, in minutes.

|  | L | LP<sub>1</sub> | LP<sub>2</sub> | LS<sub>1</sub> | LS<sub>2</sub> | LP<sub>1</sub>S<sub>1</sub> | LP<sub>2</sub>S<sub>1</sub> | LP<sub>2</sub>S<sub>2</sub> |
| --- | --- | --- | --- | --- | --- | --- | --- | --- |
| Set 2.5 mm iso | 6:22 | 8:26 | 10:29 | 8:26 | 10:29 | 10:30 | 12:33 | 14:36 |
| Set 2.0 mm iso | 8:11 | 10:50 | 13:29 | 10:50 | 13:29 | 13:29 | 16:08 | 18:47 |

To get a comparative view of the performances of protocol LS_2_, the typical multi-shell multi-tissue protocol L_msmt_ was also acquired on the second volunteer, at a spatial resolution of 2.0 mm isotropic. Since the encoding shape has a major impact on gradient waveforms optimization, leading to longer TE for STE then for LTE or PTE, this protocol was tested with two different echo times. Thus, the TE took values of 119.5 ms (same as LS_2_) and 86 ms, while keeping the same TR = 7.2 and other parameters, meaning that protocol L_msmt_ has the same acquisition time as protocol LS_2_. Furthermore, the spatial resolutions and echo times chosen for the *in vivo* acquisitions justify the SNR values described in section 3.4. Indeed, the difference in SNR between the 2.5 mm isotropic and 2.0 mm isotropic acquisitions should be given by a ratio of approximately 2, as 2.5^3^*/*2^3^ = 1.95, meaning that simulations with SNR=30 and SNR=15 can be used to study the change in resolution. As for changes in TE, simulations with SNR=15 and SNR=20 are suited to compare TE=119.5 ms with TE=86 ms.

T_1_-weighted images were also acquired at a resolution of 0.8 × 0.8 × 0.8 mm^3^ to help identifying the tissue types for the computation of the response functions.

### 3.7. Processing

The *in vivo* DW data was preprocessed using an adapted version of the Tractoflow pipeline [Theaud et al., 2020]. More precisely, the pipeline performs the MP-PCA de-noising technique (Mrtrix3) [Veraart et al., 2016], followed by susceptibility-induced distortion correction (FSL topup) [Andersson et al., 2003; Smith et al., 2004] and N4 bias field correction (ANTs) [Avants et al., 2009], on all DW images. Motion correction was not necessary on these less than 15 min long acquisitions.

The T_1_-weighted images were also treated by the pipeline, starting with a brain extraction tool (BET) [Smith, 2002] from FSL and a non-local means denoising (DIPY) [Coupé et al., 2008]. Then, the structural images were segmented into three tissues, WM, GM and CSF, using the FAST tool [Zhang et al., 2001] from FSL and finally registered on the DW images with ANTs.

This tissue segmentation led to binary masks necessary to extract axial and radial diffusivities for each tissue using a DTI model fitted with a weighted least squares method. The memsmt-CSD method was computed with a maximal SH order of 8 and with the Descoteaux07 [Descoteaux et al., 2007] SH basis adapted by DIPY, and using the default parameters of OSQP. As for the fit of the inverse Laplace transform of the gamma distribution function, the set of parameters were, following the nomenclature of [Nilsson et al., 2018]: do weight=True, do pa weight=True, do multiple s0=True, fit iters=1, guess iters=50 and all other default parameters from the authors.

## 4. Results

### 4.1. Simulated data

Figure 3 shows an example of the memsmt-CSD, DTI fit and DIVIDE outputs from the fictional anatomy described in section 3.4, using simulated protocol LP_2_S_2_ with a crossing of 90 degrees, without noise and with SNR=15. More precisely, it displays the fODFs obtained from memsmt-CSD on top of the derived volume fractions. The VF are represented with an RGB code, where red, green and blue are the CSF, GM and WM channels, respectively. For each voxel, the VF is normalized by the maximum value of all voxels, meaning that the VF keeps track of the absolute amplitude of each channel. A voxel containing partial volumes will therefore appear as a darker mix of the implicated channel, as seen in voxel 3 with the darker blue color (50% WM and 50% GM). Note that all VF voxels in this work are displayed with an opacity factor of 0.5 as well, with the purpose of increasing the fODFs visibility. Figure 3 also shows a comparison between the FA calculated from DTI and the μFA extracted with the DIVIDE method, with values comprised between 0 (black) to 1 (white).

**Figure 3:**
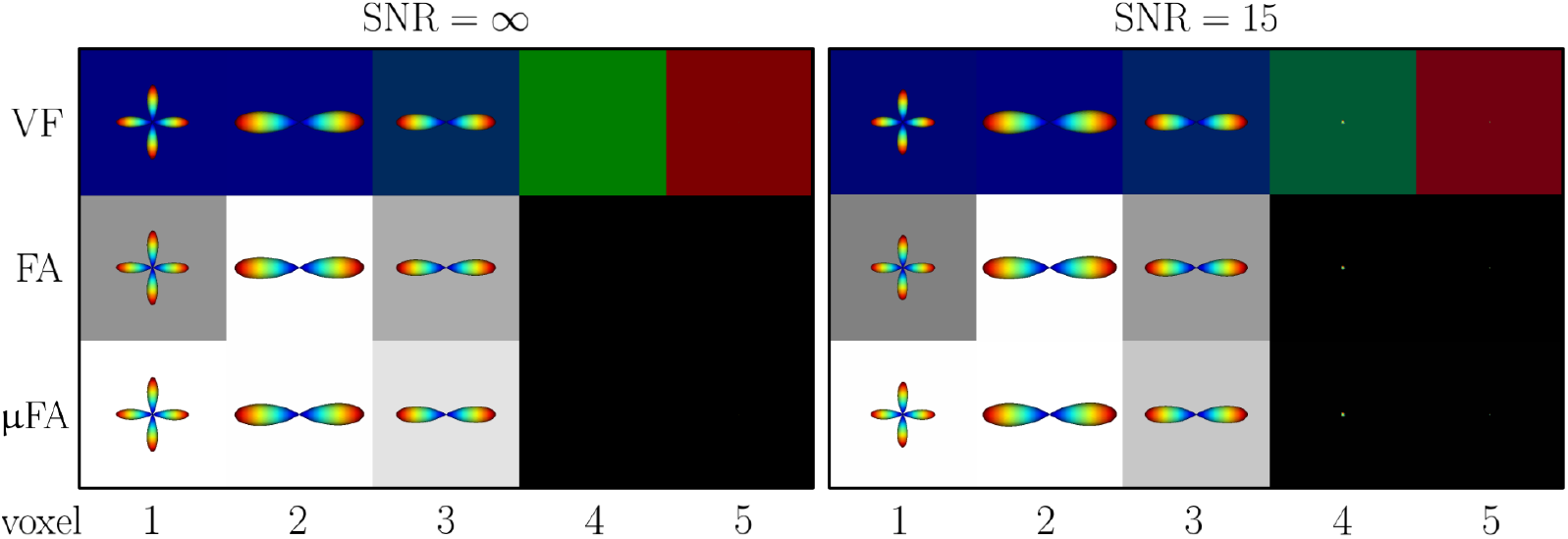
Demonstration of memsmt-CSD, DTI fit and DIVIDE methods on the simulated anatomy described in section 3.4, with protocol LP_2_S_2_ and a separation angle of 90 degrees in voxel 1, for SNR= and 15. The first row is the memsmt-CSD output, namely the WM fODFs and the volume fractions, RGB coded. The second and third rows show the FA calculated from a DTI fit and the μFA computed from DIVIDE, respectively. The fODFs are added on top of them to emphasize the contents of the voxels. FA and μFA values go from 0 to 1 (black to white color gradient).

The crossing WM fibers separation performances of each protocol from table 4 and protocol L_msmt_ are shown on figures 4, 5 and 6 for SNR=∞, SNR=30 and SNR=15, respectively. The WM fODFs are the mean of 1000 repetitions of the simulation for SNR=15 and SNR=30, and the variance of this fODFs distribution is visible as the white lobes around the fODFs, corresponding precisely to the fODFs times two standard deviations. Without noise, all protocols show a very similar performance, being able to separate the two fibers up to an angle of 55 degrees and failing for 50 degrees. The same outcome is visible at SNR=30.

**Figure 4:**
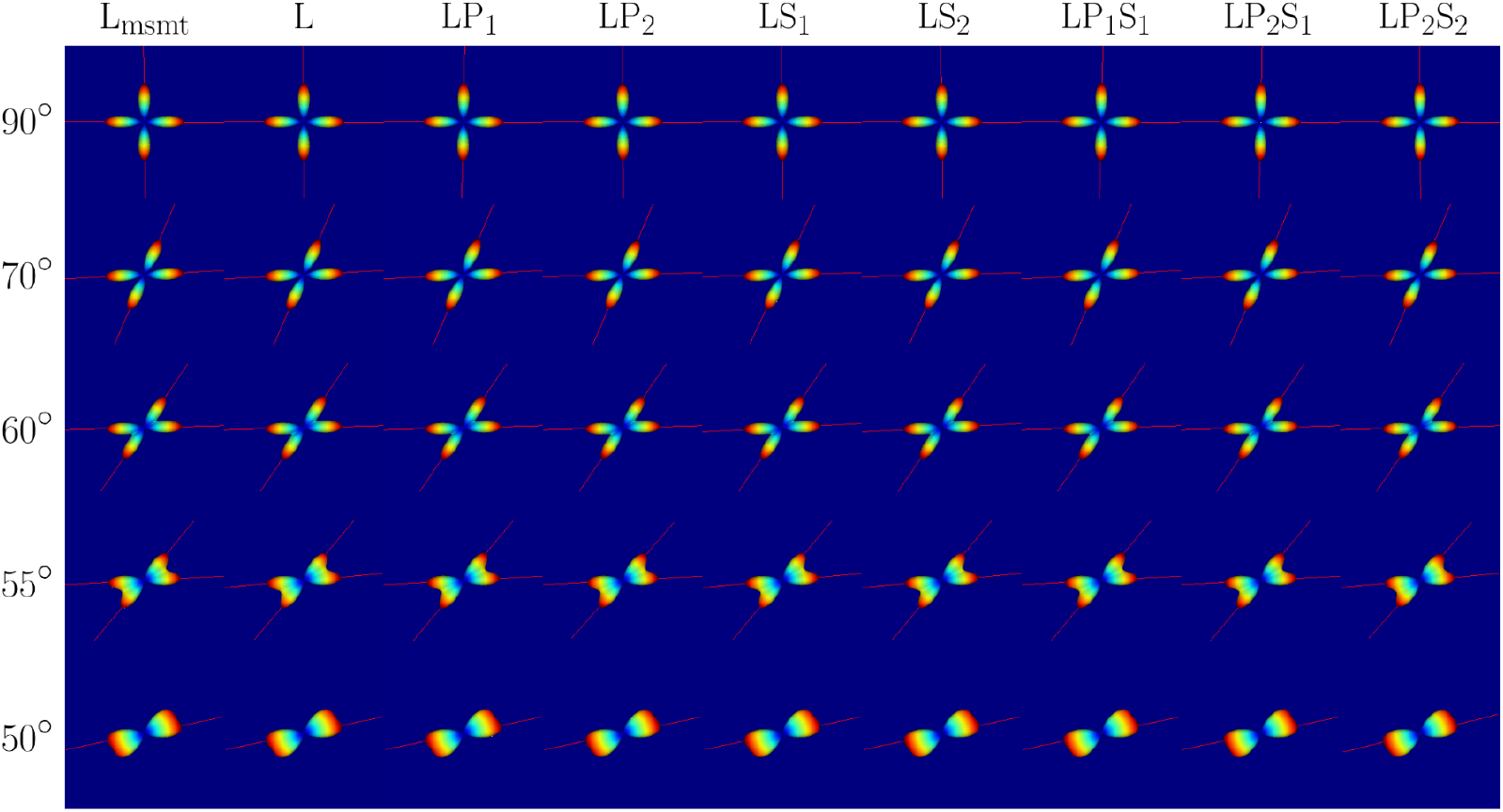
Crossing fibers of voxel 1 reconstructed with memsmt-CSD from the protocols presented in table 4 and protocol L_msmt_, at various separation angles and for SNR = ∞. In the background is the associated VF map, which is 100% WM and thus blue in this case. The red lines are the detected maxima of the fODFs.

**Figure 5:**
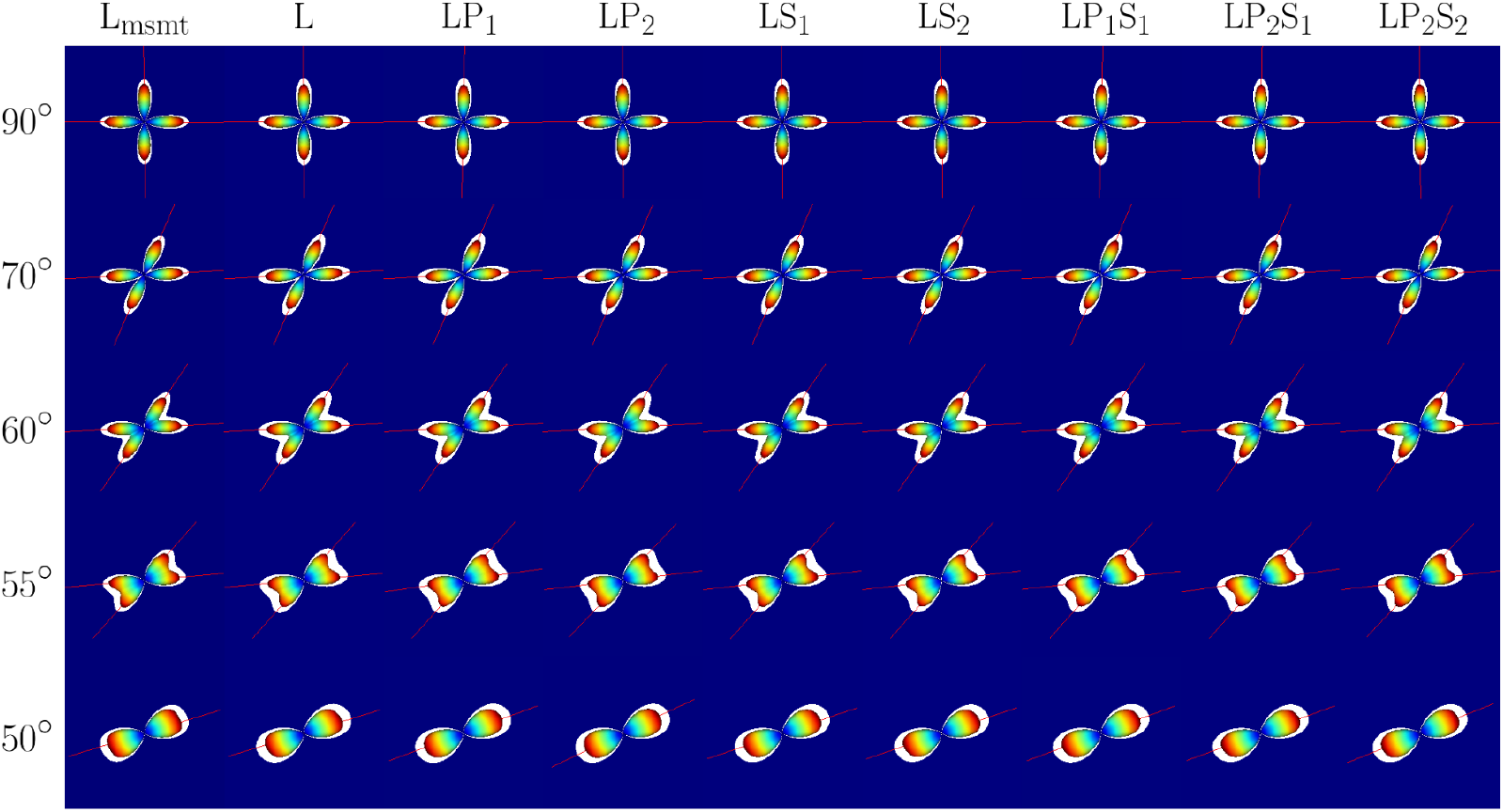
Crossing fibers of voxel 1 reconstructed with memsmt-CSD from the protocols presented in table 4 and protocol L_msmt_, at various separation angles and for SNR = 30. In the background is the associated VF map, which is 100% WM and thus blue in this case. The colored surfaces correspond to the mean fODF over 1000 simulations, whereas the white surfaces represent the mean plus two standard deviations. The red lines are the detected maxima of the fODFs.

**Figure 6:**
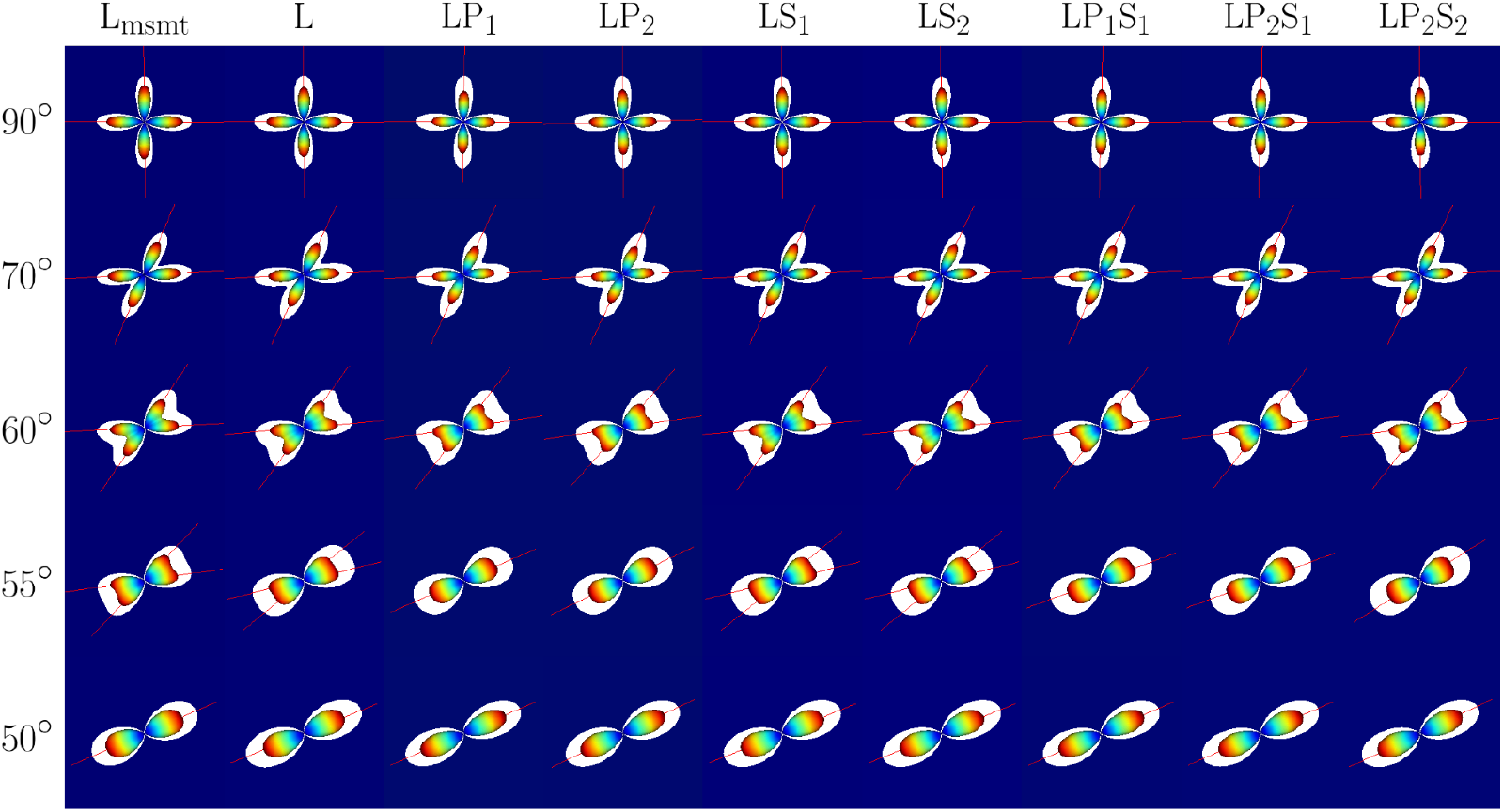
Crossing fibers of voxel 1 reconstructed with memsmt-CSD from the protocols presented in table 4 and protocol L_msmt_, at various separation angles and for SNR = 15. In the background is the associated VF map, which is 100% WM and thus blue in this case. The colored surfaces correspond to the mean fODF over 1000 simulations, whereas the white surfaces represent the mean plus two standard deviations. The red lines are the detected maxima of the fODFs.

As for SNR=15, each protocol is able to separate the two peaks up to an angle of 60 degrees, but protocols containing planar tensor encoding, P_1_ or P_2_, fail for tighter angles. The linear tensor only protocol (L), the typical multi-shell multi-tissue protocol L_msmt_ and the combined linear and spherical tensor encodings protocols (LS_1_ and LS_2_) achieve peaks separation at 55 degrees. Furthermore, these protocols seem to produce tighter fODFs, especially L_msmt_. Each protocol performs better with less noise in terms of angular resolution and precision, according to the diminishing variance of the fODFs.

Figure 7 shows a more precise representation of the angular resolution of the tested protocols at every studied SNR values, through a plot of the number of fiber orientations (NuFO) extracted with respect to the separation angle. This points out the exact angle at which the protocols fail to separate the two fibers, noticeable by the drops in the curves, allowing a better understanding of the behaviours presented on figures 4, 5 and 6.

**Figure 7:**
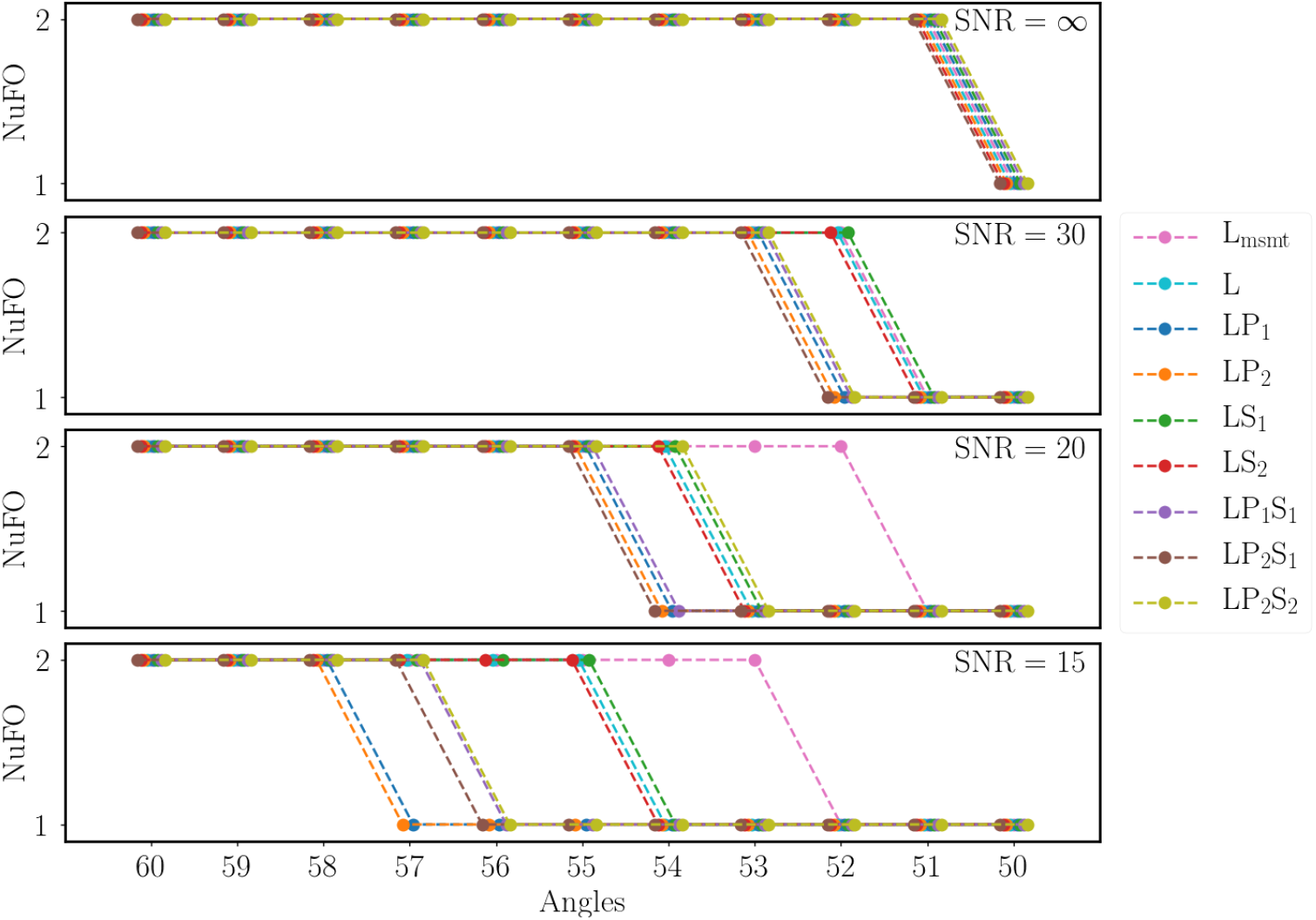
Angular dependency of the number of fiber orientations extracted from the mean fODF over 1000 simulations with memsmt-CSD from the studied protocols in voxel 1, at various SNR. Note that for each angle, the dots are slightly shifted to better distinguish the protocols, but touching dots are truly at the same angle value.

Without noise, all protocols perform the same, losing the WM fiber crossing right before the 50 degrees mark, at 51 degrees, as the NuFO drops from 2 to 1. At SNR=30, protocols L, L_msmt_, LS_2_ and LS_1_ start to separate themselves from protocols containing planar tensor encoding (P_1_ or P_2_), as they lose 1 degree from SNR=∞, whereas all the other protocols lose 2 degrees. Protocol L_msmt_ remains the same at SNR=20, distancing itself from protocols L, LS_2_ and LS_1_, which drop from 52 to 54 degrees of angular resolution. At this SNR level, all protocols with PTE also drop by 2 degrees (55 degrees), except for LP_2_S_2_, which has an angular resolution of 54 degrees. While figure 6 shows a seemingly big difference (5 degrees) between L, L_msmt_, LS_2_ and LS_1_ and the rest of the protocols at SNR=15, figure 7 indicates that all protocols differ from a maximum of 3 degrees at this SNR, with the exception of L_msmt_. Indeed, the combination of only linear and planar encodings (LP_2_ and LP_1_) has the lowest angular resolution at 58 degrees, followed closely by the protocols composed of all encoding types (LP_1_S_1_, LP_2_S_1_ and LP_2_S_2_) with 57 degrees. The L, LS_2_ and LS_1_ protocols are able to distinguish tighter angles, dropping from 2 to 1 NuFO after 55 degrees. The L_msmt_ protocol has the highest angular resolution with 53 degrees. Note that protocol LP_2_ was also tested using the L gradient table instead of the L** gradient table to make sure that the directions redistribution does not impact the angular resolution of memsmt-CSD. Since the results are exactly the same as the normal LP_2_, the points are not shown on figure 7.

The precision and accuracy of the DIVIDE method at computing the μFA with different protocols is presented on figure 8, where the μFA from every protocol, as well as the ground truth μFA, are plotted for the five simulated voxels, for each SNR. Throughout each protocol, the μFA computed without noise is following the ground truth value, while the noisy data deviates from it. The error bars, computed from the variance over the 1000 repetitions, also show the precision of the protocols. It is worth noting that the μFA computed for every protocol at each voxel is always overlapping with the ground truth value when considering the error bars, for both SNR=15, 20 and 30. For voxels 1 to 3 at these SNR values, results of protocols LS_2_, LP_2_S_1_ and LP_2_S_2_ are significatively different from those of other protocols (p-values < 0.01), while being mostly similar between themself (p-values > 0.01), according to the p-values calculated from t-tests. These protocols are consistently more precise and also show better accuracy, being closer to the ground truth most of the time in voxels 1 to 3. As for voxels 4 and 5, all protocols with noisy data produce approximately the same overestimated μFA and are not statistically different (p-values > 0.01). Still for SNR=15, 20 and 30, protocols LP_2_, LS_1_ and LP_1_S_1_ all perform similarly for each voxel (p-values > 0.01), while protocol LP_1_ has by far the worst accuracy and precision, especially in voxels 1 to 3. However, it is important to notice that protocol LP_2_ at SNR=20 is still statistically different then protocols LS_2_, LP_2_S_1_ and LP_2_S_2_ at SNR=15. Furthermore, its precision and accuracy at SNR=20 are also lower then protocols LS_2_, LP_2_S_1_ and LP_2_S_2_ at SNR=15, even though they are much closer.

**Figure 8:**
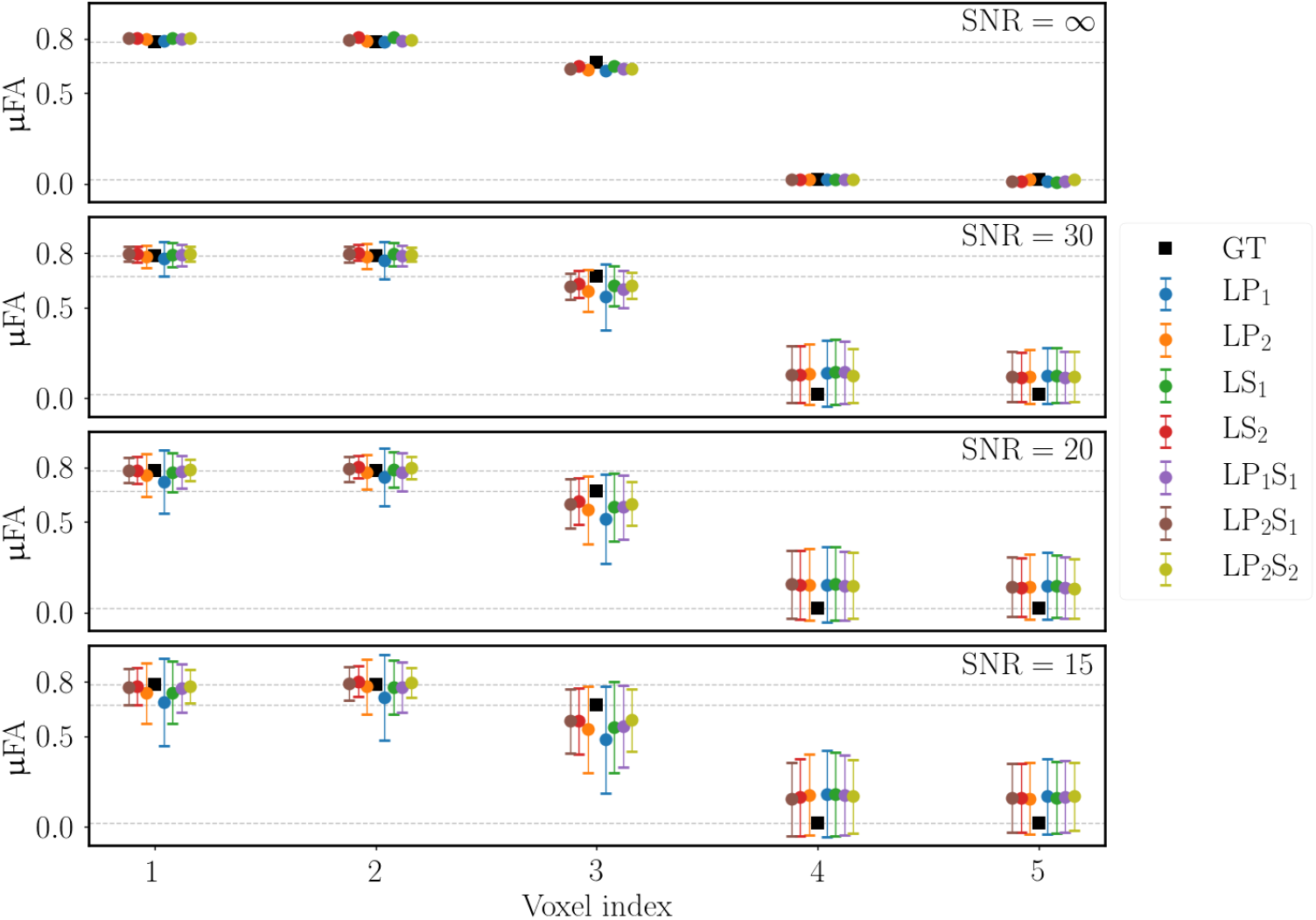
Mean μFA over 1000 simulations for each voxel and each protocols, at various SNR. The error bars correspond to the standard deviation extracted from the 1000 repetitions of the simulations. The black squares and the horizontal dashed grey lines represent the ground truth μFA.

### 4.2. In vivo data

Figures 9 and 11 show memsmt-CSD outputs with protocol LS_2_ on *in vivo* data acquired with resolutions of 2.5 × 2.5 × 2.5 mm^3^ and 2 × 2 × 2 mm^3^ respectively, following the preprocessing steps discussed in section 3.7 (see figure A.15 for examples of raw data). Once again, the RGB coded volume fractions allow to distinguish between the three studied tissues, WM, GM and CSF. More precisely, these figures display a zoom of chosen brain slice that exhibit voxels similar to the fictional anatomy studied by simulation. Indeed, figures 9 and 11 contain pure single WM fiber and WM fiber crossing voxels, as well as CSF and GM voxels. In the top right of these figures, an image of the whole brain slice helps locating the zoomed image.

**Figure 9:**
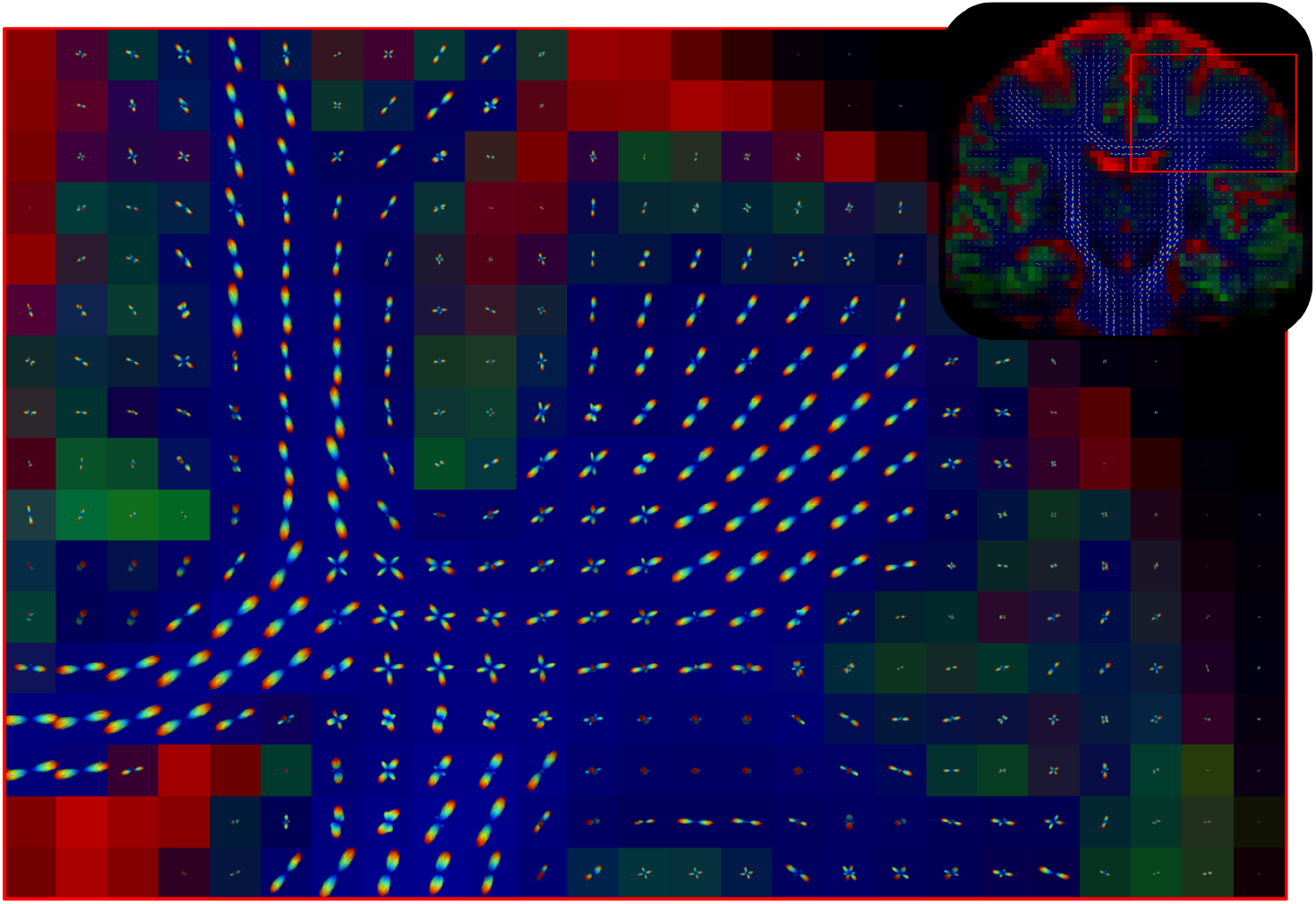
Section of an *in vivo* human brain coronal slice at 2.5 mm isotropic showing the WM fODFs obtained from memsmt-CSD with protocol LS_2_, on top of the computed volume fractions map. The VF is RGB coded, with red being the CSF, green the GM and blue the WM.

**Figure 10:**
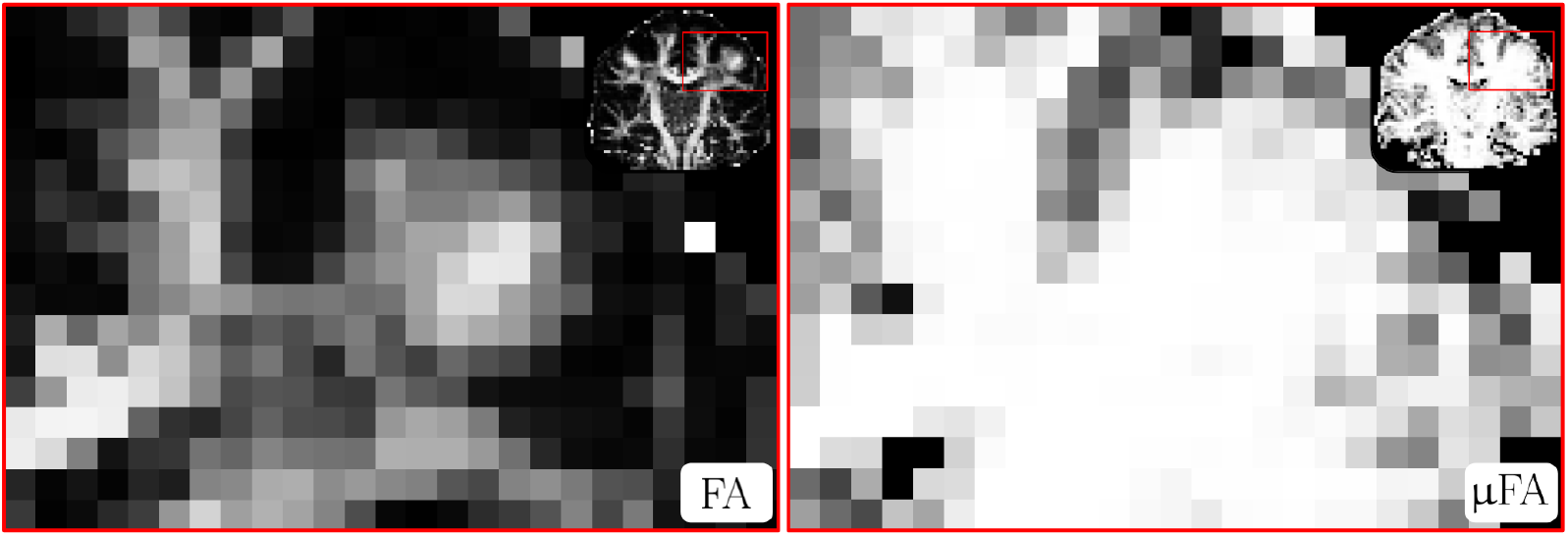
Same brain coronal slice as figure 9, still at 2.5 mm isotropic, showing the FA computed from DTI with protocol L and the μFA computed from DIVIDE with protocol LS_2_. Both measures have values going from 0 (black) to 1 (white).

**Figure 11:**
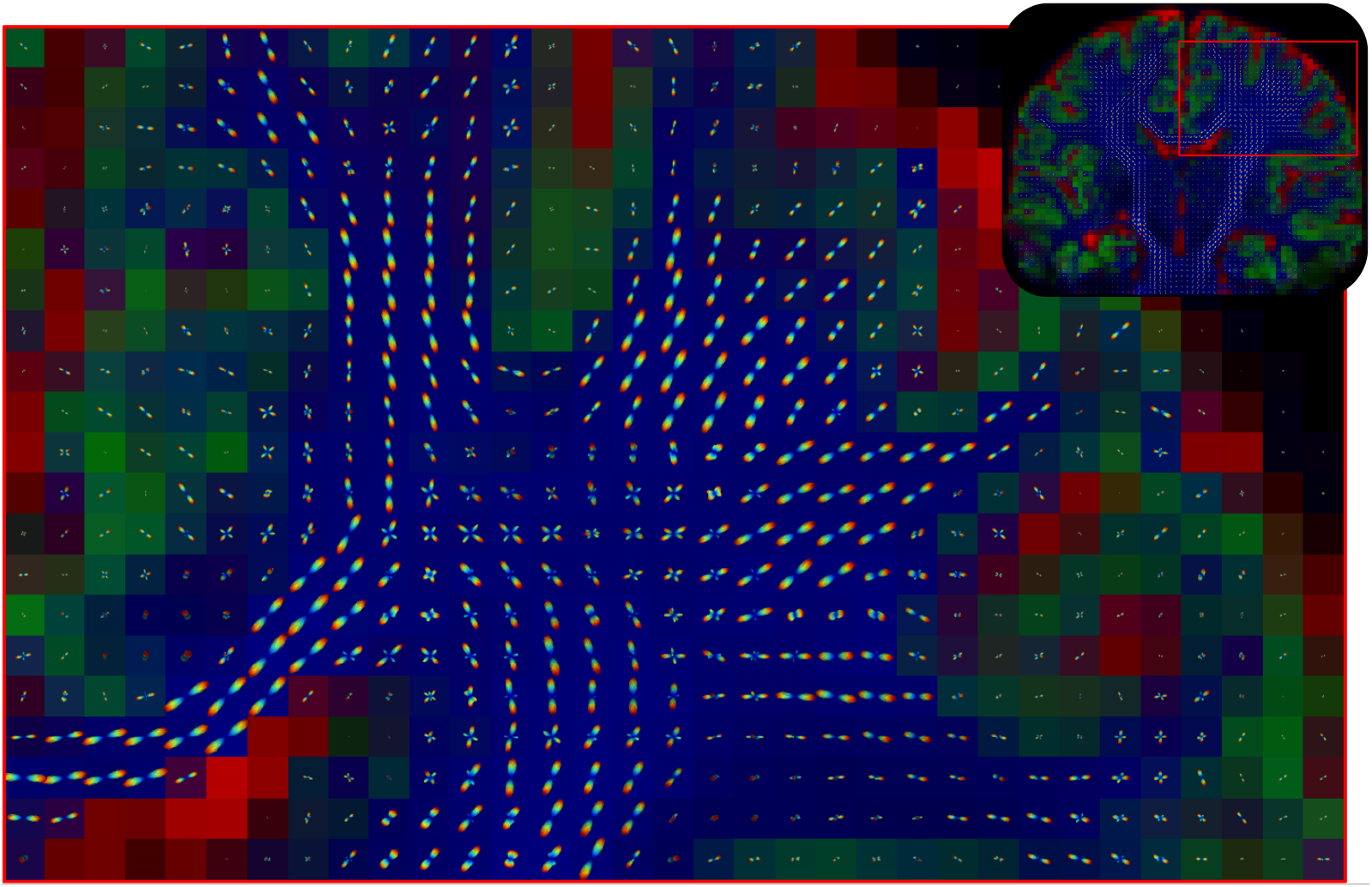
Section of an *in vivo* human brain coronal slice at 2.0 mm isotropic showing the WM fODFs obtained from memsmt-CSD with protocol LS_2_, on top of the computed volume fractions map. The VF is RGB coded, with red being the CSF, green the GM and blue the WM.

Figures 10 and 12 present the FA and μFA for the same brain slice as figures 9 and 11, respectively. The FA, calculated only from the LTE protocol (L), shows high intensity voxels where single WM fibers are present, and lower intensity where WM fibers are crossing (see figures 9 and 11 for comparison with fODFs). The μFA in the other hand, calculated from the combination of LTE and STE for protocol LS_2_, shows high intensity voxels for all of the WM voxels, regardless of the presence of crossings.

**Figure 12:**
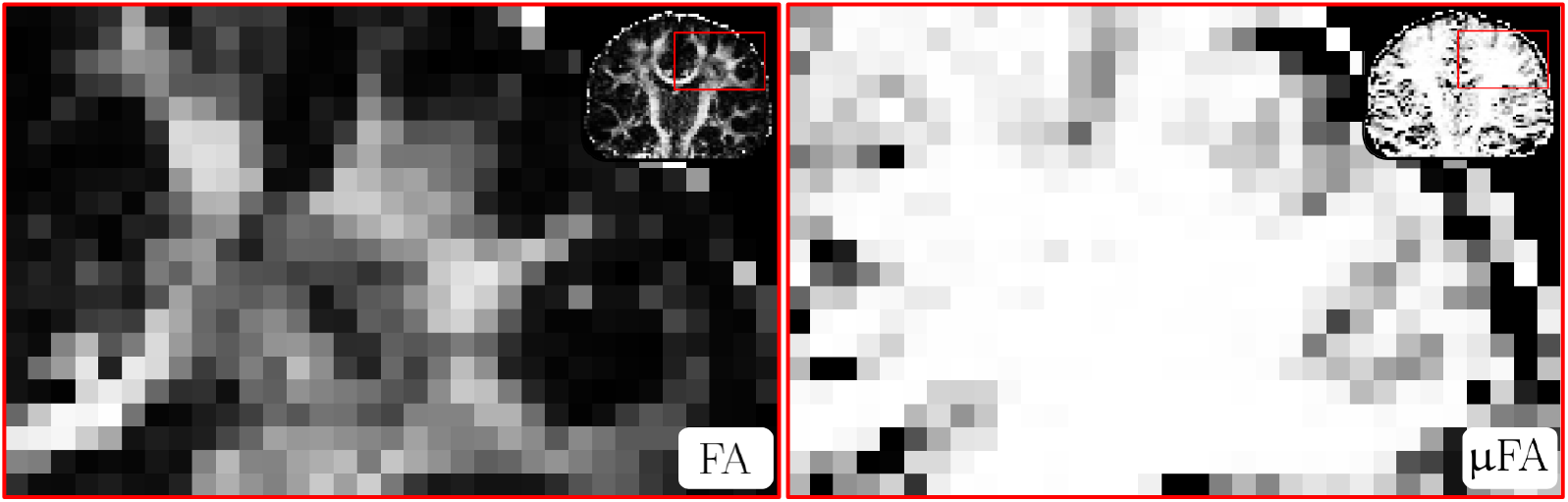
Same brain coronal slice as figure 11, still at 2.0 mm isotropic, showing the FA computed from DTI with protocol L and the μFA computed from DIVIDE with protocol LS_2_. Both measures have values going from 0 (black) to 1 (white).

Figure 13 presents msmt-CSD outputs obtained with the short echo time (86 ms) version of protocol L_msmt_ on *in vivo* data acquired with a resolution of 2 × 2 × 2 mm^3^.

**Figure 13:**
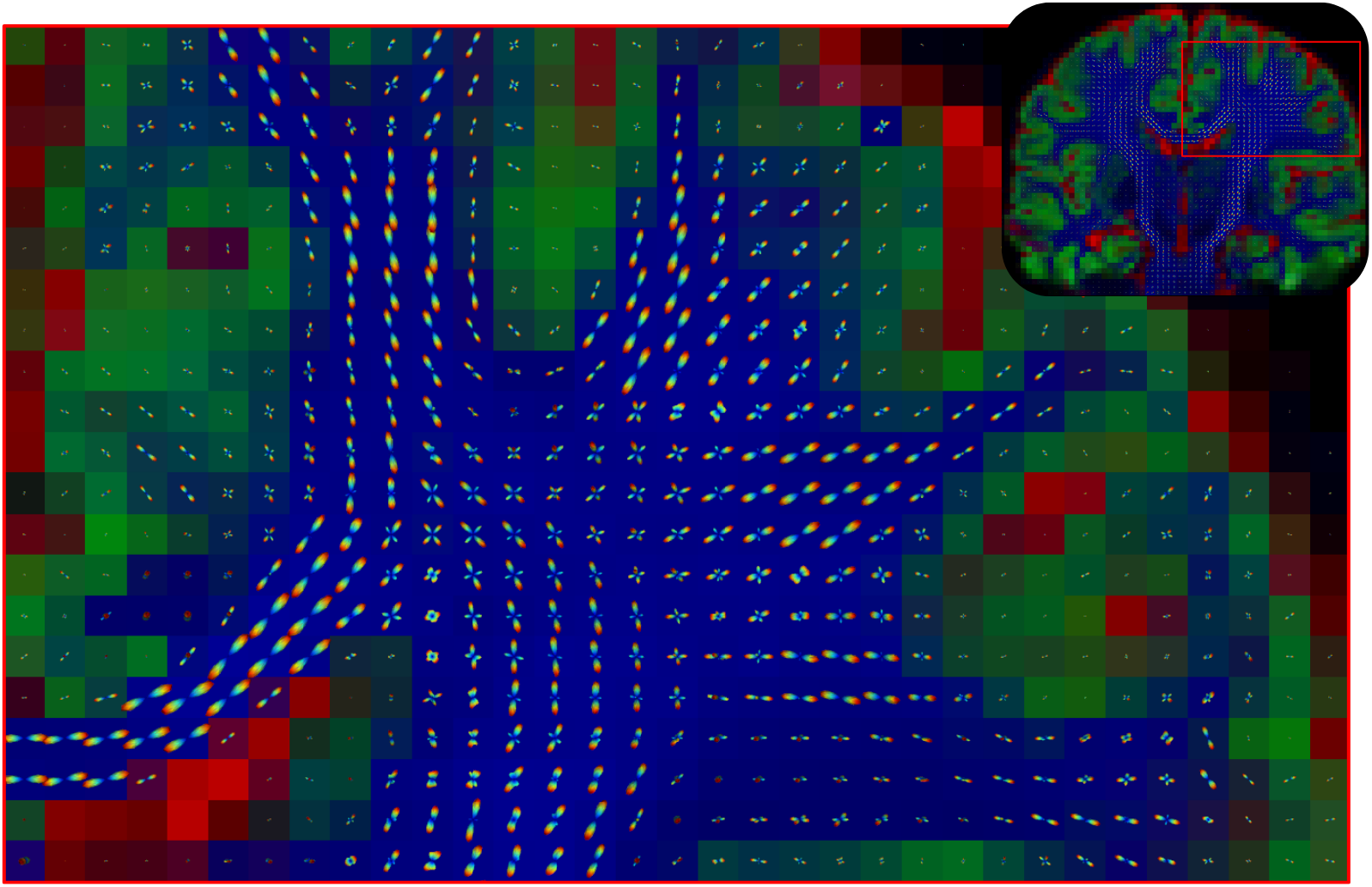
Section of an *in vivo* human brain coronal slice at 2.0 mm isotropic showing the WM fODFs obtained from msmt-CSD with the short TE version of protocol L_msmt_, on top of the computed volume fractions map. The VF is RGB coded, with red being the CSF, green the GM and blue the WM.

## 5. Discussion

### 5.1. Choices of b-values and number of directions per shell

The choices of b-values and number of gradient directions per shell, previously presented in table 3, were inspired by [Nilsson et al., 2020]. In this paper, the authors present a three-minute tensor-valued dMRI protocol enabling the calculation of the diffusional variances, and thus, the μFA, via a powder-averaged two-term cumulant expansion approach. This approach is comparable to the DIVIDE method used in our study, according to Reymbaut et al. [2020c]. The adaptations made from Nilsson et al. [2020] affect both the b-values and the number of directions per shells. Their protocol consisted on four b-values (*b* = 100, 700, 1400, and 2000 s/mm^2^) acquired in 3, 3, 6, and 6 directions for the linear encoding, and 6, 6, 10, and 16 directions for the spherical encoding. We modified the b-values to get a *b* = 1200 s/mm^2^ for the DTI fit [Jones and Basser, 2004], swapping it with the *b* = 1400 s/mm^2^. We also diminished the *b* = 2000 s/mm^2^ to a *b* = 1800 s/mm^2^ to keep a smaller gap between the b-values to help fit equation 6, and we added a *b* = 2400 s/mm^2^ shell for CSD purposes. As shown by figure 8, this distribution of b-values seems to provide sufficient coverage to get proper μFA values.

The number of directions per shell taken from Nilsson et al. [2020] ensures a good enough directional coverage for the rotation invariant criteria. However, this number is too low to perform CSD, or even to get a well defined DTI fit. Based on the results from Jones [2004], we chose to take 12 directions for the *b* = 1200 s/mm^2^ shell in linear encoding. Figure 3 indeed shows that the FA calculated from DTI with this configuration is in agreement with the literature [Pierpaoli and Basser, 1996], with high values in voxels of single WM fiber (voxel 2), lower values in WM fiber crossings (voxel 1) and near-zero values in voxels with only isotropic contents (voxel 4 and 5). This figure also justifies the choice of 12 directions by the fact that the FA does not change much from data without noise to data of SNR=15. Note that the DTI fit can also be achieved on planar and spherical encodings, allowing the calculation of FA and MD with planar encoding and MD with spherical encoding. However, we chose to use only the linear encoding for DTI measures computation, as fits from other encodings seemed to be heavily sensitive to noise. As for the two higher b-values, 18 and 24 directions were taken, respectively, for a total of 54 directions over *b* = 1000 s/mm^2^ available for the standard msmt-CSD with linear encoding. The performances of protocol L for SNR=15, presented on figures 6 and 7, confirm that this amount of directions, combined with the lower shells, is enough to propel the angular resolution up to 55 degrees with msmt-CSD.

Since the proposed number of directions for the spherical encoded gradients S_2_ mostly serves for SNR purposes and not for directional coverage, as a single spherical encoded signal is already rotation invariant [Szczepankiewicz, 2016], we reduced the number of directions to 3, 3, 6 and 6, creating a spherical encoding gradient table S_1_ with shorter acquisition time, at the price of lower SNR. Figure 8 show that for low SNR (15), the DIVIDE method is indeed affected by the decrease of STE directions, as LS_1_ is significantly less precise than LS_2_. As for memsmt-CSD, it does not seem to be impacted by this effect, as demonstrated by figure 7, where both LS_1_ and LS_2_ perform as well as the LTE only protocol (L). Therefore, reducing the number of directions from S_2_ to S_1_ might not be good for μFA computation, but it does not affect the fODFs computation.

In addition to linear and spherical encodings, two planar encoding gradient tables P_1_ and P_2_ were created with the same amount of directions that were tested for linear encoding by Nilsson et al. [2020]. The smaller one, P_1_, should be sufficient to get rotation invariant signals, since planar encoding requires less directions than linear encoding in that matter [Szczepankiewicz, 2016; Szczepankiewicz et al., 2019]. Considering this, the results from figure 8, that show that protocol LP_1_ is the worst in terms of accuracy and precision when it comes to μFA computation with noise, even when comparing at SNR=20 to take into account a possible lower TE due to the absence of STE, probably indicate that planar tensor encoding suffers more from noise than spherical tensor encoding in terms of DIVIDE. Indeed, even with more directions (LP2) at SNR=20, PTE still does not produce results as good as protocol LS_2_ at SNR=15, which has the same amount of directions.

### 5.2. Effects of protocol choice on the computation of μFA

The choice of protocol has a big impact on the accuracy and the precision of the computed μFA, as shown by figure 8. According to this figure, PTE alone with LTE (LP_1_ and LP_2_) is consistently underperforming other protocols in voxels containing WM (voxels 1 to 3) and when comparing at equal SNR. In these voxels, we demonstrate that STE and LTE combined can produce accurate and precise measures of μFA, if a sufficient amount of directions is used, which is the case of LS_2_. This observation agrees with the conclusions of Nilsson et al. [2020] about their proposed protocol, very similar to LS_2_. Furthermore, the use of three tensor shapes with enough directions can also provide good μFA measures, as LP_2_S_1_ and LP_2_S_2_ display accuracy and precision very similar to LS_2_. However, the higher acquisition times needed for LP_2_S_1_ and LP_2_S_2_ do not procure significant advantage to the computation of the μFA, leaving LS_2_ with the best time to quality ratio. Moreover, when taking into account the fact that STE requires higher TE then LTE or PTE and thus suffers from lower SNR values, the use of a protocol such as LP_2_ becomes interesting. Nevertheless, figure 8 demonstrates that even when comparing LP_2_ at SNR=20 with LS_2_ at SNR=15, which mimics the effect of various echo times similar to the ones used in the *in vivo* acquisitions, the LS_2_ protocol is still producing slightly better results then LP_2_.

In voxels 4 and 5, composed of GM and CSF respectively, a constant overestimation of the μFA is observed. This is due to the noise inducing apparent anisotropy at the microscopic scales, increasing the μFA. Indeed, this is confirmed by the fact that this effect appears only when adding noise, since μFA is accurately estimated at SNR=∞ and starts to deviate more and more with added noise. In fact, this increase of anisotropy measures with noise is a well known phenomenon discussed in Pierpaoli and Basser [1996]; Jones and Basser [2004] and caused by eigenvalue repulsion [Mehta, 2004]. Ultimately, no protocol stands out as the better one for these two voxels.

### 5.3. Effects of protocol choice on the angular resolution of a WM fiber crossing

The choice of protocol has a smaller impact for the results of memsmt-CSD than it has for the DIVIDE method. Indeed, figures 4, 5 and 6 present the effects of adding planar and/or spherical tensor shapes to a LTE protocol. Instinctively, it would be surprising to see a complete breakdown when adding additional tensor-valued encodings at a constant SNR level. As a matter of fact, figure 7 shows that adding a STE gradient table (S_1_ or S_2_) to the linear (L) protocol does not diminish the minimal angle at which CSD is able to distinguish two WM fibers crossing. Nevertheless, the figure also shows that adding either of the two PTE gradient tables (P_1_ or P_2_) to the linear (L) protocol decreases the angular resolution of the CSD method by 3 degrees at SNR=15 and by 1 degree at SNR=20. This effect is indeed more visible at low SNR, meaning that PTE might not react well with noise when it comes to distinguishing two WM fiber populations. Adding STE to LP_1_ or LP_2_ seems to lower this negative effect, as it allows memsmt-CSD to reach 1 more degree of separation at SNR=15. Once again, to provide with a fairer comparison, protocols containing STE should be compared at SNR=15 with other protocols at SNR=20, to emulate the effect of a shorter TE. When doing so, it appears that protocols LS_1_ and LS_2_ indeed have the same angular resolution as protocols LP_1_ and LP_2_ (55 degrees) and that these protocols all have 1 less degree of angular resolution then protocol L. Furthermore, it is worth noting that the typical multi-shell protocol with the same amount of gradient directions as protocols LS_2_ or LP_2_ has 3 degrees more of angular resolution at SNR=20. However, protocol L_msmt_ does not provide μFA measures with DIVIDE. If the acquisition time is brought to the discussion (see table 5), three tensor shapes protocols (LP_1_S_1_, LP_2_S_1_ and LP_2_S_2_) are not worth the longer time, and protocols LS_1_, LS_2_, LP_1_ and LP_2_ all stand out as the better choices of multi-encoding protocols for computing WM fibers crossing with memsmt-CSD.

### 5.4. Combined memsmt-CSD and DIVIDE performances

While figure 3 serves as a proof of concept for the memsmt-CSD, DTI fit and DIVIDE methods, figures 7 and 8 bring to light the importance of the chosen protocol for fODFs angular resolution and μFA computation. Indeed, when combining the conclusions of section 5.2 and 5.3, protocol LS_2_ arises as the best choice for computing both memsmt-CSD and DIVIDE. Even if the typical L_msmt_ protocol shows the better results for CSD, reorganising the gradient directions distribution to create LS_2_ from the same amount of directions only leads to the lost of a few degrees of angular resolution, while enabling the DIVIDE process. Still, protocol LP_2_ must not be forgotten, as it has proven to be comparable to LS_2_ in terms of angular resolution with memsmt-CSD and to produce μFA measures that are only slightly less precise and accurate then protocol LS_2_.

Figures 9 and 10 provide an example of what protocol LS_2_ can achieve in terms of WM fODF reconstruction and measures calculation at a typical resolution for tensor-valued dMRI data (2.5 mm isotropic). Figures 11 and 12 show that this resolution can be increased to 2.0 mm isotropic and still produce valuable WM fODF reconstruction and measures calculation. Figures 9 and 11 show that memsmt-CSD, combined with protocol LS_2_, is able to produce clean WM fODFs in WM voxels, but most importantly in crossing fibers voxels. Furthermore, it is able to distinguish WM from GM and CSF, rendering a sort of tissue segmentation. Figures 10 and 12 show that the LTE part of protocol LS_2_ can be used to produce a FA map that represent very well the WM voxel, when compared to the WM segmentation from memsmt-CSD, apart from the famous FA drop in crossing fibers voxels. Besides, figures 10 and 12 also show that the same protocol can be used to compute a μFA map, which does not suffer like the FA in crossing fibers. However, the μFA seems to be swelled a bit, especially in regions of WM and GM interface. This phenomena is probably attributable to the noise level of these *in vivo* acquisition, but perhaps a better calibration of the DIVIDE method could be done. Nevertheless, those figures prove that memsmt-CSD is viable and that fODFs, FA and μFA can all be computed from protocol LS_2_.

Moreover, figure 13 allows to compare the performances of msmt-CSD and memsmt-CSD, using protocol L_msmt_ with a lower echo time. It appears that both methods produce similar fODFs and volume fractions map separating the different tissues. However, it is also clear that msmt-CSD and memsmt-CSD do not reconstruct the same results. Indeed, the GM parts of the VF map seem to be different and the fODFs in WM crossing also show differences.

### 5.5. Recommendations and future work

Considering all of the above, we recommend using our combination of LTE and STE gradient tables to create the LS_2_ protocol in the scope of computing both fODFs and μFA. Indeed, this protocol uses two different tensor encoding shapes for an accurate and precise computation of the μFA with a DIVIDE method, while not losing too much angular resolution with memsmt-CSD in comparison to similar msmt-CSD. Furthermore, its acquisition time of approximately 10 minutes at a resolution of 2.5 mm isotropic makes it attractive for research purposes. If possible, we recommend spending 3 more minutes to push the resolution up to 2.0 mm isotropic and get less partial volume voxels. As structural connectivity, tractometry and connectomics is becoming more and more present in research and disease applications, it is important to push for higher spatial resolution multi-dimensional acquisition in clinically feasible times.

While the *in vivo* results of memsmt-CSD are promising, the computed μFA could benefit from more denoising or better DIVIDE tuning. Nevertheless, the proposed protocol is a starting point for future research, where fODFs and μFA are both of interest. Further investigation of the impact of multiple b-tensor shapes on the VF values and fODF reconstruction in voxels of partial volume would be needed. Indeed, the interesting problem of the presence of an isotropic compartment inside a WM fiber crossing voxel could be better suited for a protocol containing STE, like LS_2_. Moreover, the impact of planar tensor encoding on memsmt-CSD should be explored in more depth and is not to be forgotten. Further investigation of the pros and cons of memsmt-CSD, especially with STE, could provide a better understanding of the effects of this technique on microstructure estimation at the voxel level but also at the tractography and connectome level. More digging and a better comparison with msmt-CSD should be done, especially regarding the difference in the VF maps.

## 6. Conclusion

In this work, we first established the mathematical and computational foundations of a multi-encoding msmt-CSD model, able to compute fODFs and volume fraction maps from tensor-valued dMRI data. Using simulated data, we showed that the model can indeed produce multi-tissue volume fraction maps and white matter fODFs in single and crossing fibers voxels. Furthermore, these fODFs only suffer from the lost of a few degrees in terms of angular resolution when adding sufficient amount of spherical tensor encoding or planar tensor encoding acquisitions to a linear tensor encoding. Moreover, the performance of different combinations of linear, planar and spherical gradient tables were also evaluated with the DIVIDE method on the simulated data. We showed that while combining three b-tensor shapes provides great accuracy and precision in μFA computation, combining only linear and spherical b-tensors also yields competing accuracy and precision values, while producing satisfying fODF reconstruction with memst-CSD.

We thus propose a 10 min protocol at 2.5 mm isotropic and a 13 min protocol at 2 mm isotropic combining linear and spherical b-tensor encodings to get both memsmt-CSD and diffusional variance decomposition methods. These protocols therefore kill two birds with one stone, allowing the reconstruction of accurate crossing fiber fODFs for tractography/connectivity while being able to extract the μFA map precisely.

## 7. Acknowledgements

The authors wish to thank François Rheault and Guillaume Theaud for their help with the implementation of the memsmt-CSD method in DIPY, as well as Charles Poirier for his help with fODF visualization with variances. They are also grateful to the Fonds de recherche du Québec - Nature et technologies (FRQNT) and the Natural Sciences and Engineering Research Council of Canada (NSERC) programs for funding this research. The authors also thank Prof. Christine Tardif and Prof. Pascal Tétreault for their advice on this paper.

They also wish to thank the HCP project for the use of their data for estimating the non-DW signals per tissues. The HCP project (Principal Investigators: Bruce Rosen, M.D., Ph.D., Martinos Center at Massachusetts General Hospital; Arthur W. Toga, Ph.D., University of Southern California, Van J. Weeden, MD, Martinos Center at Massachusetts General Hospital) is supported by the National Institute of Dental and Craniofacial Research (NIDCR), the National Institute of Mental Health (NIMH) and the National Institute of Neurological Disorders and Stroke (NINDS). Collectively, the HCP is the result of efforts of co-investigators from the University of Southern California, Martinos Center for Biomedical Imaging at Massachusetts General Hospital (MGH), Washington University, and the University of Minnesota.

## Appendix A. Datasets & Databases

**Figure A.14:**
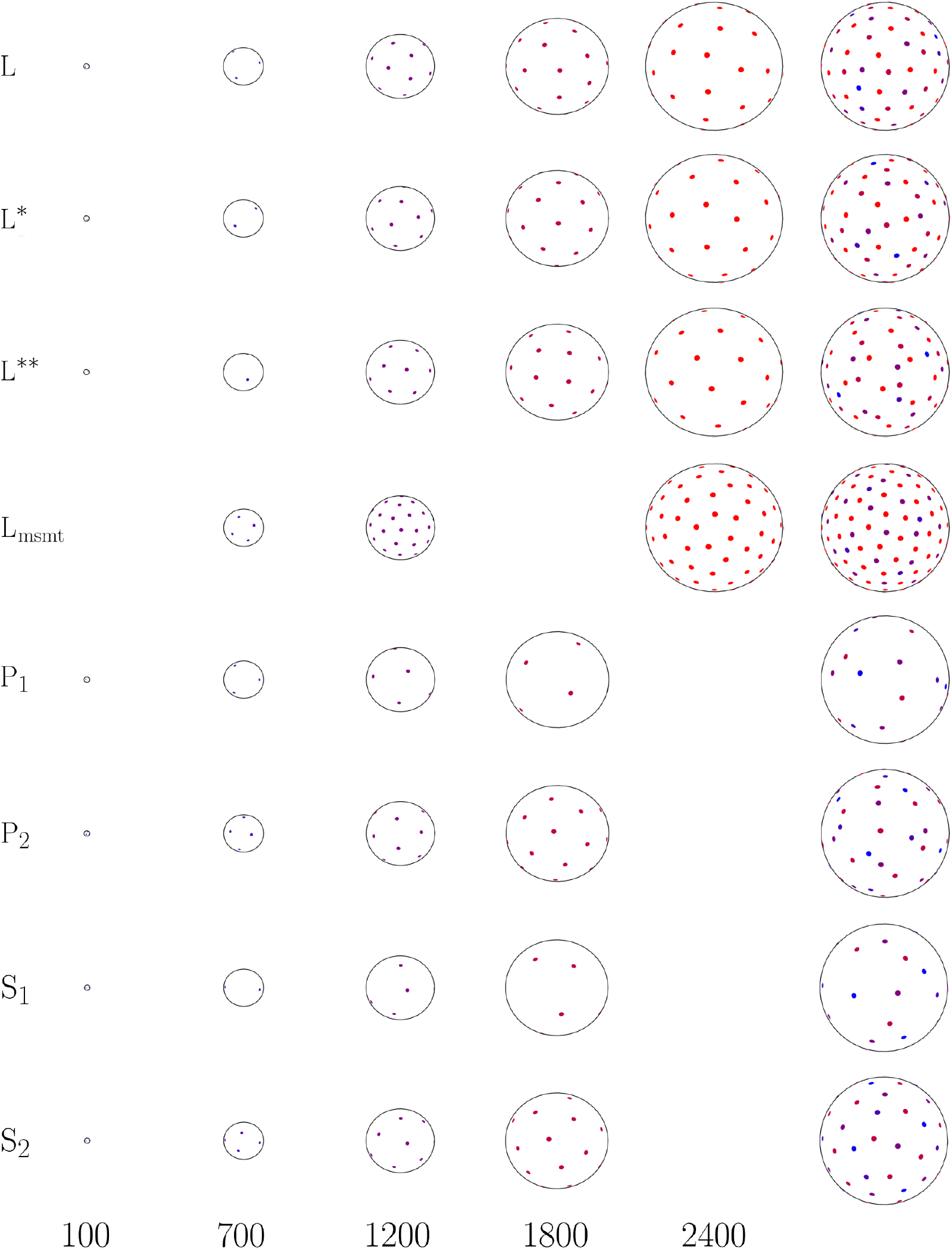
Visualization of the gradient directions distribution for each protocols studied. The first five columns represent the distribution at b-values of 100, 700, 1200, 1800 and 2400 s/mm^2^. For these columns, the amplitudes of the spheres are proportional to the b-values. The last column is every shell combined on one sphere. The exact gradient tables are available at https://doi.org/10.5281/zenodo.4628539.

**Figure A.15:**
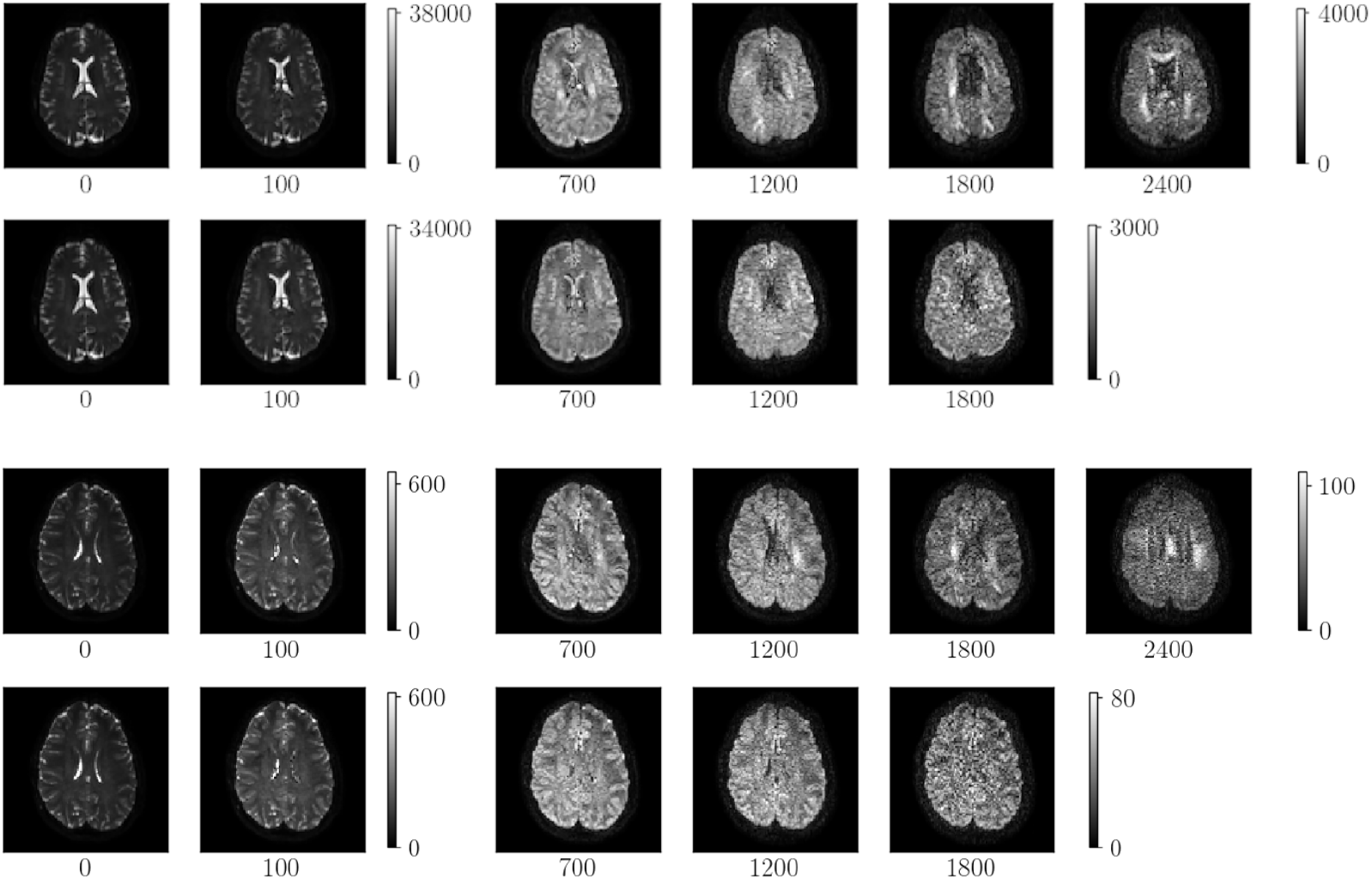
Examples of the raw data used in this study. The two first rows are images from the 2.5 mm isotropic dataset, with the first of these row being the L gradient table and the second one being S_2_. The two last rows are images from the 2.0 mm isotropic dataset, with the first of these row being the L gradient table and the second one being S_2_. Under each image is the b-value, in s/mm^2^.

